# The Causal Effects of Education on Health, Mortality, Cognition, Well-being, and Income in the UK Biobank

**DOI:** 10.1101/074815

**Authors:** Neil M Davies, Matt Dickson, George Davey Smith, Gerard van den Berg, Frank Windmeijer

## Abstract

Educated people are generally healthier, have fewer comorbidities and live longer than people with less education. Previous evidence about the effects of education come from observational studies many of which are affected by residual confounding. Legal changes to the minimum school leave age is a potential natural experiment which provides a potentially more robust source of evidence about the effects of schooling. Previous studies have exploited this natural experiment using population-level administrative data to investigate mortality, and relatively small surveys to investigate the effect on mortality. Here, we add to the evidence using data from a large sample from the UK Biobank. We exploit the raising of the school-leaving age in the UK in September 1972 as a natural experiment and regression discontinuity and instrumental variable estimators to identify the causal effects of staying on in school. Remaining in school was positively associated with 23 of 25 outcomes. After accounting for multiple hypothesis testing, we found evidence of causal effects on twelve outcomes, however, the associations of schooling and intelligence, smoking, and alcohol consumption may be due to genomic and socioeconomic confounding factors. Education affects some, but not all health and socioeconomic outcomes. Differences between educated and less educated people may be partially due to residual genetic and socioeconomic confounding.

**Significance Statement:** On average people who choose to stay in education for longer are healthier, wealthier, and live longer. We investigated the causal effects of education on health, income, and well-being later in life. This is the largest study of its kind to date and it has objective clinic measures of morbidity and aging. We found evidence that people who were forced to remain in school had higher wages and lower mortality. However, there was little evidence of an effect on intelligence later in life. Furthermore, estimates of the effects of education using conventionally adjusted regression analysis are likely to suffer from genomic confounding. In conclusion, education affects some, but not all health outcomes later in life.

**Funding:** The Medical Research Council (MRC) and the University of Bristol fund the MRC Integrative Epidemiology Unit [MC_UU_12013/1, MC_UU_12013/9]. NMD is supported by the Economics and Social Research Council (ESRC) via a Future Research Leaders Fellowship [ES/N000757/1]. The research described in this paper was specifically funded by a grant from the Economics and Social Research Council for Transformative Social Science. No funding body has influenced data collection, analysis or its interpretations. This publication is the work of the authors, who serve as the guarantors for the contents of this paper. This work was carried out using the computational facilities of the Advanced Computing Research Centre -http://www.bris.ac.uk/acrc/ and the Research Data Storage Facility of the University of Bristol — http://www.bris.ac.uk/acrc/storage/. This research was conducted using the UK Biobank Resource.

**Data access:** The statistical code used to produce these results can be accessed here: (https://github.com/nmdavies/UKbiobankROSLA). The final analysis dataset used in this study is archived with UK Biobank, which can be accessed by contacting UK Biobank.

## Introduction

On average educated people are healthier, happier, more intelligent, richer, and live longer than those with less education. We do not know whether this is because education directly causes these outcomes, by affecting behaviors, such as smoking, or if these differences are due to other factors, such as socioeconomic or genomic differences. Whether education causes differences in outcomes later in life has been the subject of significant debate by epidemiologists, economists and other social scientists.(1–14) Economists have argued that a substantial portion of the benefits of education accrue via its potential effects on mortality and morbidity.(14) Epidemiologists have found that people who attended university have higher fluid intelligence in adulthood.(15) These associations are robust to adjustment for parental social class and adolescent cognition, which has been taken by some as proof that education causes later outcomes.(16) Despite this, many epidemiologists and economists are acutely aware that correlation, and multivariable adjusted regression, can be unreliable evidence of causation.(17–19) The ideal study design to definitively prove whether education has causal effects would be to randomize the age at which children leave school. However, this experiment would not be ethical, cost-effective, or timely. A more feasible, but potentially robust, research design is to exploit natural experiments that affected when people left school but are not related to confounding factors.(20, 21) One widely used form of natural experiment is to exploit changes in the legal minimum school leaving age. These changes forced some people to stay in school for longer than they would have otherwise chosen.

In September 1972, the United Kingdom raised the school-leaving age from age 15 to 16. Researchers have previously used this policy change to investigate the effects of forcing students to stay in school longer using administrative data and longitudinal cohort studies.(1, 22–24) However, the cohort studies had relatively small samples and, as a result, had to include people who were born many years before and after the reform. These studies produced relatively imprecise estimates of the effects of education. Previous results from administrative data lacked detailed information on covariates to identify people born in England affected by the reform or measures of many of the outcomes of interest such as cognition.

In the current study, we used the raising of the school-leaving age in 1972 as a natural experiment to estimate the causal effects of schooling. We used a regression discontinuity design and used novel data from the UK Biobank.(25, 26) We add to the literature in two ways. First, this is the largest sample with detailed individual-level information from the school years immediately before and after the reform. Second, we used genome-wide data to prove that the observational associations of education and other outcomes are likely to suffer from genomic confounding.

## Results

Of the 502,644 participants in the UK Biobank, who were all aged between 37 and 73 at recruitment in 2008, 423,106 were born in England, see Figure 1 for a flow diagram of exclusion and inclusion of participants in this study. See Table 1 for a description of their characteristics. The youngest participants, those born between 1960 and 1971 obtained more education than those born earlier in the twentieth century (Figure 2). This is consistent with the well-documented secular increase in the length of education over the period.(1) UK Biobank includes 11,240 and 10,898 participants who turned 15 years old in the first year before and the first year after the school-leaving age increased. Before the reform, 85% of participants stayed in school after the age of 15, whereas after the reform almost 100% of participants remained in school after the age of 15. The proportions of men and women who remained in school after age 15 increased over time (Figures 3 and 4). Participants born in July and August could still technically leave school before their 16^th^ birthday, so on average participants born in the summer term report leaving school at a younger age.

**Figure 1:**
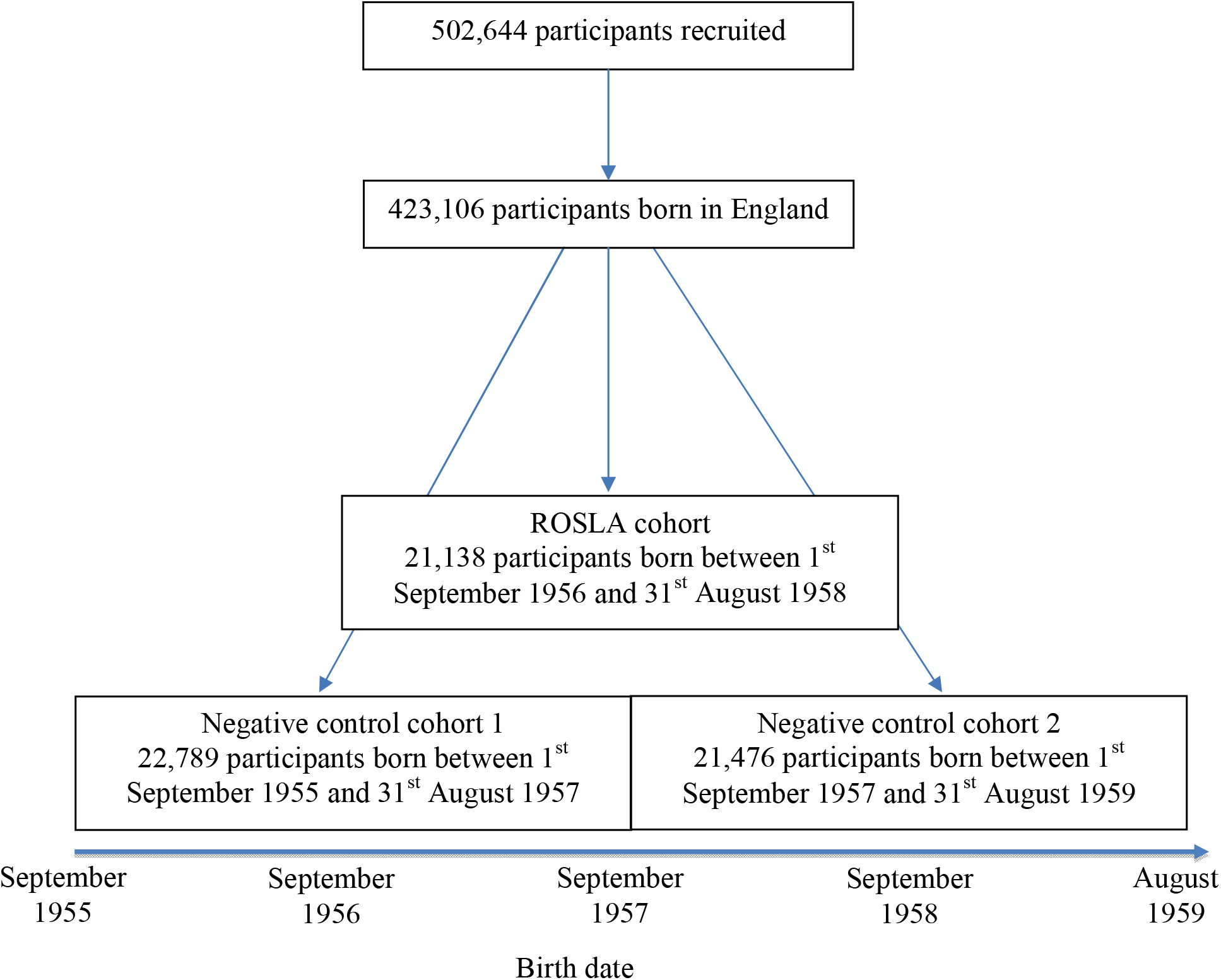
Flow chart of inclusion and exclusion of participants into the study.

**Figure 2:**
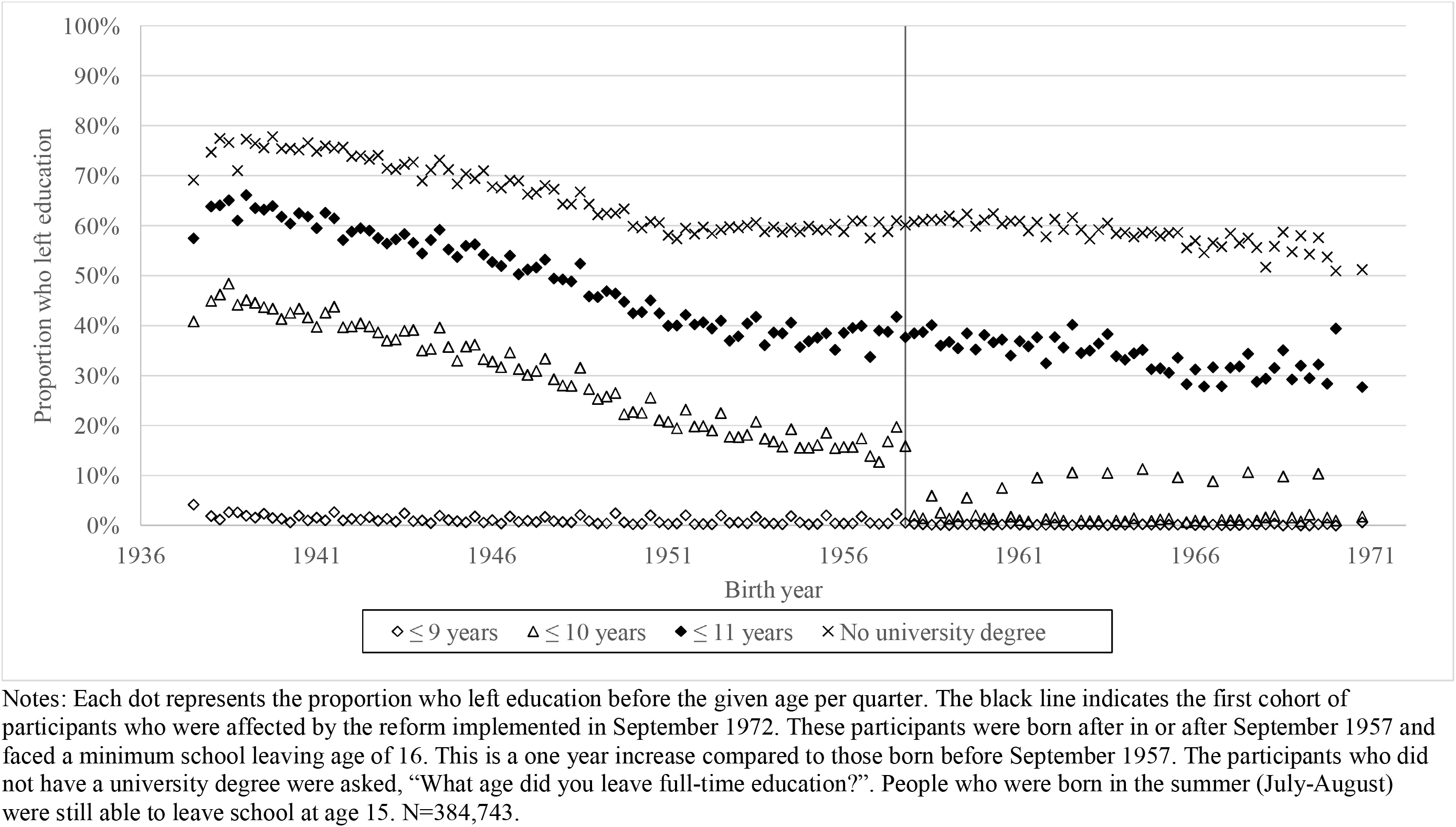
Years of full-time education by quarter of birth.

**Figure 3:**
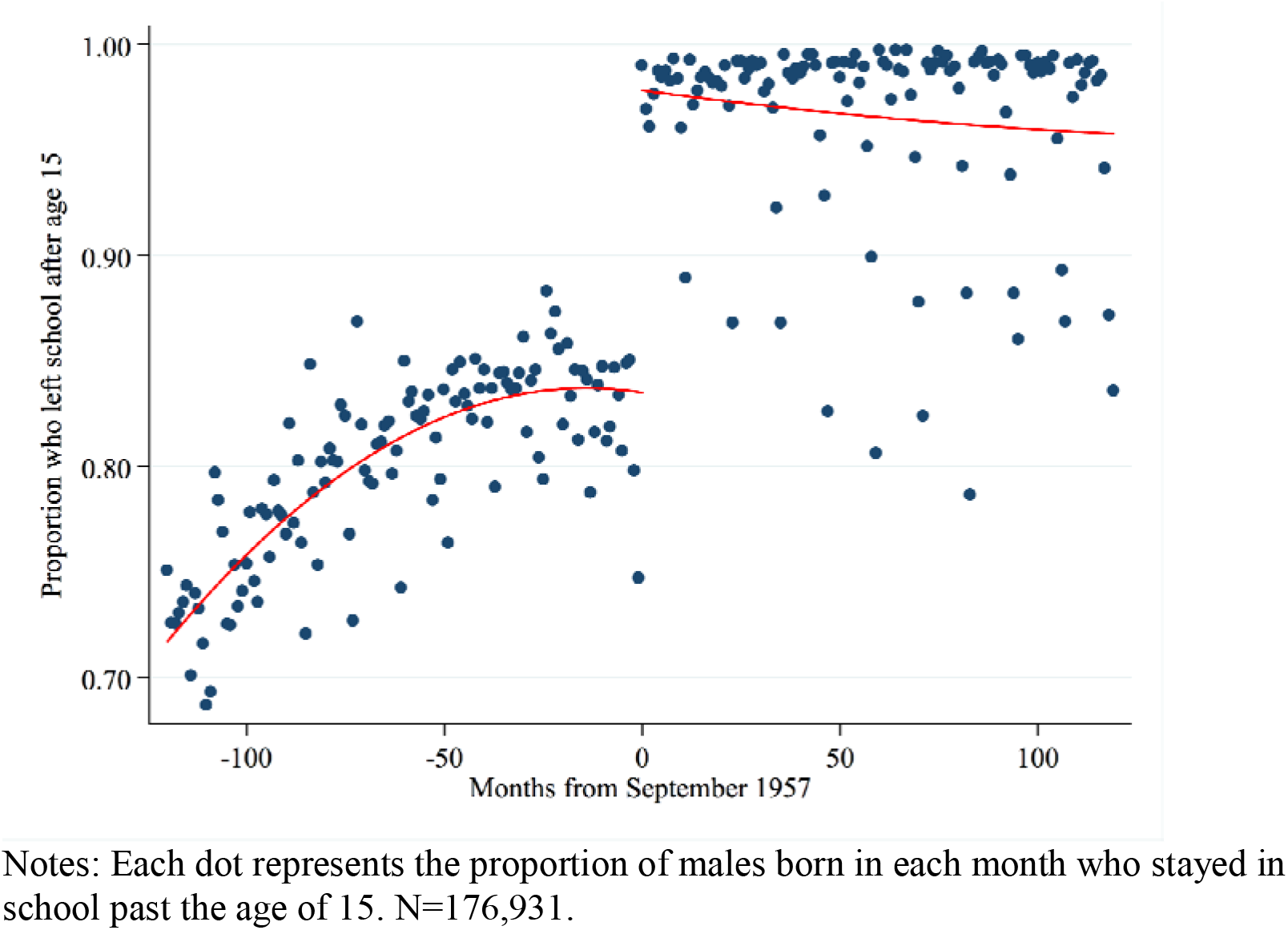
Effects of the reform on the proportion of males staying in education after the age of 15.

**Figure 4:**
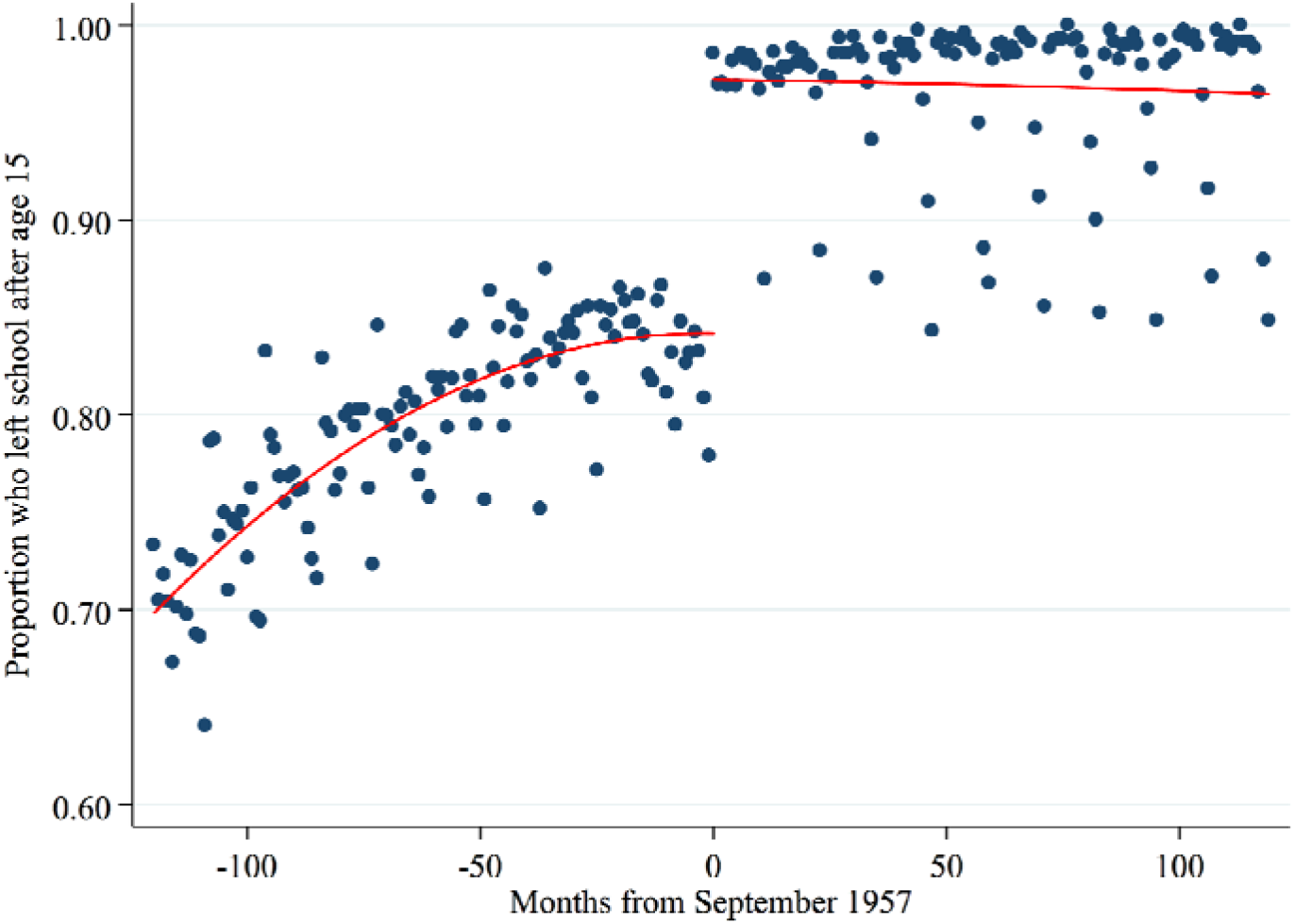
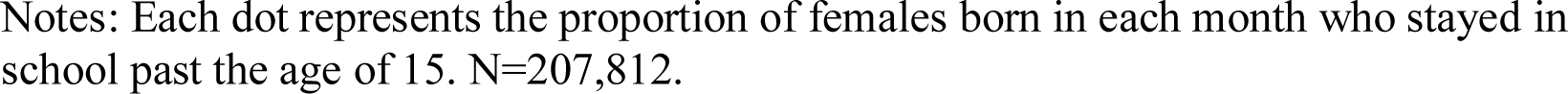
Effects of the reform on the proportion of females staying in education after the age of 15.

**Table 1:** Cohort of UK Biobank participants born between September 1956 and August 1958.

|  | N | Count | Proportion (%) |  |  |
| --- | --- | --- | --- | --- | --- |
| Male | 22,138 | 9,699 | 43.8 |  |  |
| Mother smoked during pregnancy | 19,442 | 6,519 | 33.5 |  |  |
| Breastfed | 18,226 | 12,695 | 69.7 |  |  |
| Father alive | 21,618 | 8,198 | 37.9 |  |  |
| Mother alive | 21,822 | 13,388 | 61.4 |  |  |
| High blood pressure | 21,768 | 3,978 | 18.3 |  |  |
| Diabetes | 22,049 | 661 | 3.0 |  |  |
| Stroke | 22,110 | 184 | 0.8 |  |  |
| Heart attack | 22,110 | 183 | 0.8 |  |  |
| Cancer | 22,011 | 1,813 | 8.2 |  |  |
| Died | 22,138 | 191 | 0.9 |  |  |
| Ever smoker | 22,086 | 8,899 | 40.3 |  |  |
| Current smoker | 22,086 | 2,602 | 11.8 |  |  |
| Income over £18k | 19,921 | 17,398 | 87.3 |  |  |
| Income over £31k | 19,921 | 13,532 | 67.9 |  |  |
| Income over £52k | 19,921 | 7,524 | 37.8 |  |  |
| Income over £100k | 19,921 | 1,638 | 8.2 |  |  |
|  | N | Mean | Standard deviation | Minimum | Maximum |
| Birthweight (kg) | 14,860 | 3.32 | 0.63 | 0.57 | 7.26 |
| Number of brothers | 21,848 | 1.15 | 1.18 | 0.00 | 12.00 |
| Number of sisters | 21,851 | 1.06 | 1.13 | 0.00 | 14.00 |
| Genome-wide allele education score | 7,005 | -0.01 | 0.97 | -4.37 | 3.48 |
| Grip strength (kg) | 21,989 | 1.49 | 10.85 | -30.66 | 46.46 |
| Arterial Stiffness | 8,537 | -0.32 | 3.04 | -8.34 | 83.63 |
| Height (cm) | 22,077 | 169.42 | 9.16 | 122.00 | 206.00 |
| BMI (kg/m<sup>2</sup>) | 22,055 | 27.32 | 4.96 | 14.53 | 61.54 |
| Diastolic blood pressure (mmHg) | 21,494 | 82.55 | 10.29 | 45.00 | 131.50 |
| Systolic blood pressure (mmHg) | 21,492 | 133.50 | 16.82 | 84.00 | 268.00 |
| Intelligence (0 to 13) | 8,540 | 6.34 | 2.10 | 0.00 | 13.00 |
| Happiness (0 to 5 Likert) | 8,626 | 3.36 | 0.72 | 0.00 | 5.00 |
| Alcohol consumption (1 low, 5 high) | 22,123 | 3.14 | 1.42 | 0.00 | 5.00 |
| Hours watching television per day | 21,206 | 2.63 | 1.59 | 0.00 | 24.00 |
| Moderate exercise (days/week) | 21,330 | 3.46 | 2.34 | 0.00 | 7.00 |
| Vigorous exercise (days/week) | 21,379 | 1.91 | 1.95 | 0.00 | 7.00 |

### Covariate Balance Tests

People who stayed in school after age 15 had higher birth weights; mothers who were less likely to smoke during pregnancy; were taller than average at age 10 relative to their peers; were more likely to have been breastfed; had fewer siblings; and were more likely to have parents who were alive; they were more likely to have SNPs known to be associated with higher educational attainment(27) (Table 2). In comparison, there were few detectable baseline differences between people affected and unaffected by the reform. The only detectable difference was that participants in the first year affected by the reform were 4.3 (95% confidence intervals (95%CI): 2.5 to 6.1) and 3.7 (95%CI: 2.6 to 4.8) percentage points more likely to have an alive father and mother when they attended the assessment center in 2008-2010. There was weak evidence that fewer participants in the younger cohort were breastfed. The participants affected by the reform are, by definition, one year younger than those who were not affected. The raw differences above do not account for this age difference. We investigated whether these associations were simply due to the effects of age by estimating the effects of two negative control “placebo” reforms. For the first placebo reform, we constructed a sample of the two school years born before the reform (negative control sample 1) and for the second, we used the two first two school years after the reform (negative control sample 2). We estimated the difference in outcomes between the school years in exactly the same way as described above for the true reform. As with the true reform, on average there is a year difference in age between the two school years in the negative control samples. Therefore, if the differences between those unaffected by the reform and those affected were solely due to aging, then we should see similar differences between the two school cohorts who turned 15 before the reforms occurred, and the two school cohorts who turned 15 after the reform occurred. We know that all the participants in the two negative control samples experienced the same school leaving age. We found few differences between the two school years in negative control sample 1, or negative control sample 2. The only detectable differences were that the younger cohorts in both cases were more likely to have parents who were alive and were less likely to be breastfed (Table 3). These results suggest that participants unaffected and affected by the reform had similar observed covariates. This suggests that environmental or genomic confounding is unlikely to bias estimates which use the raising of the school-leaving age to identify the effects of education. In contrast socioeconomic and genomic factors confound the associations of school and later outcomes. Participants who *chose* to stay in school had more advantaged backgrounds and were more likely to have genetic variants associated with educational attainment.

**Table 2:** Baseline differences between students who left before and after age 15 (left) and differences between students who left school before and after the reform (right).

| Independent variable: |  | Stayed on in school after age 15 |  |  |  | ROSLA cohort vs. pre-ROSLA cohort |  |  |  |
| --- | --- | --- | --- | --- | --- | --- | --- | --- | --- |
| Dependent variable: | N | Risk | 95% Confidence interval |  | P- | Risk | 95% Confidence interval |  | P- |
|  |  | difference | Lower | Upper | value | difference | Lower | Upper | value |
| Male | 22,138 | -0.003 | -0.024 | 0.019 | 0.79 | 0.000 | -0.016 | 0.016 | 0.98 |
| Mother smoked during pregnancy | 19,442 | -0.161 | -0.185 | -0.137 | 1.5E-8 | -0.010 | -0.024 | 0.003 | 0.13 |
| Breastfed | 18,226 | 0.095 | 0.074 | 0.117 | 9.9E-7 | -0.014 | -0.028 | 0.001 | 0.07 |
| Father alive | 21,618 | 0.104 | 0.084 | 0.123 | 1.6E-7 | 0.043 | 0.025 | 0.061 | 2.9E-4 |
| Mother alive | 21,822 | 0.110 | 0.089 | 0.130 | 1.3E-7 | 0.037 | 0.026 | 0.048 | 1.2E-5 |
|  |  | Mean |  |  |  | Mean |  |  |  |
|  |  | difference |  |  |  | difference |  |  |  |
| Birthweight (kg) | 14,860 | 0.033 | 0.000 | 0.066 | 0.05 | 0.007 | -0.019 | 0.033 | 0.56 |
| Number of brothers | 21,848 | -0.507 | -0.596 | -0.419 | 6.9E-8 | -0.031 | -0.077 | 0.016 | 0.17 |
| Number of sisters | 21,851 | -0.405 | -0.518 | -0.292 | 7.4E-6 | 0.010 | -0.031 | 0.051 | 0.61 |
| Genome-wide education allele Z-score | 7,005 | 0.015 | 0.011 | 0.019 | 3.0E-6 | 0.001 | -0.001 | 0.002 | 0.22 |

Notes: ROSLA= Raising of the school leaving age. The results in the left hand columns present the association of staying on school after age 15 and the covariates in the two cohorts directly before and after the reform. The right columns present the differences in these covariates between participants who were born before immediately before and after the reform. Robust standard errors clustered by year and month of birth reported.

**Table 3:** Association of the reform in two dummy control populations. First participants who attended school in the two years prior to the reforms (negative control sample 1, left), and in the two years after the reforms (negative control sample 2, right).

| Independent variable: |  | Negative control sample 1 |  |  |  | Negative control sample 2 |  |  |  |  |
| --- | --- | --- | --- | --- | --- | --- | --- | --- | --- | --- |
| Sample: |  | Sept 1955-August 1956 to Sept 1956-August 1957 |  |  |  | Sept 1957- August 1958 to Sept 1958-August 1959 |  |  |  |  |
|  |  | Risk | 95% Confidence interval |  | P- |  | Risk | 95% Confidence interval |  | P- |
| Dependent variable: | N | difference | Lower | Upper | value | difference | difference | Lower | Upper | value |
| Male | 22,789 | 0.011 | 0.001 | 0.022 | 0.04 | 21,476 | 0.002 | -0.014 | 0.019 | 0.75 |
| Mother smoked during pregnancy | 19,992 | -0.005 | -0.016 | 0.005 | 0.29 | 18,893 | -0.013 | -0.031 | 0.005 | 0.14 |
| Breastfed | 18,674 | -0.018 | -0.033 | -0.003 | 0.02 | 17,748 | -0.016 | -0.032 | -0.001 | 0.04 |
| Father alive | 22,232 | 0.025 | 0.013 | 0.038 | 0.001 | 20,924 | 0.041 | 0.027 | 0.054 | 4.2E-5 |
| Mother alive | 22,447 | 0.032 | 0.026 | 0.039 | 2.8E-7 | 21,157 | 0.032 | 0.019 | 0.046 | 2.3E-4 |
|  |  | Mean |  |  |  |  | Mean |  |  |  |
|  |  | difference |  |  |  |  | difference |  |  |  |
| Birthweight (kg) | 14,970 | 0.016 | -0.008 | 0.040 | 0.17 | 14,591 | 0.008 | -0.009 | 0.025 | 0.33 |
| Number of brothers | 22,478 | 0.025 | -0.017 | 0.066 | 0.21 | 21,193 | 0.018 | -0.005 | 0.040 | 0.11 |
| Number of sisters | 22,478 | -0.012 | -0.036 | 0.011 | 0.28 | 21,195 | 0.015 | -0.032 | 0.062 | 0.49 |
| Genome-wide education allele Z-score | 7,134 | 0.000 | -0.003 | 0.003 | 0.88 | 6,702 | 0.000 | -0.004 | 0.003 | 0.89 |

Notes: The left hand columns compare the two cohorts immediately before the reform, and the right hand columns compare the two cohorts immediately after the reform. Robust standard errors clustered by year and the month of birth reported. Linear regression adjusted for month of birth and gender.

### Reduced form

We report two comparisons: first, the differences between participants who chose to stay in school after the age of 15 and those who left, and second, the difference between participants who were not affected by the reform (those born before September 1957) and those who were affected by it (those born in or after September 1957). On average participants who stayed in school after age 15 had better outcomes later in life. Fewer educated participants were diagnosed with high blood pressure; diabetes; a stroke; or a heart attack; or died (left columns in Table 4). There were minor differences in cancer diagnoses by education level. Fewer participants who stayed in school were diagnosed with depression. Educated people had stronger grip strength, lower arterial stiffness; and lower blood pressure. They were also taller, lighter, and achieved higher scores on intelligence tests. Participants who left school at age 15 reported similar levels of happiness as those who stayed on. Educated people drank more alcohol, but were much less likely to smoke. They were more likely to report higher incomes; watched less television, but exercised less. There were fewer differences between the participants affected and unaffected by the reform (right columns in Table 4). Participants affected by the reform were less likely to report having high blood pressure, diabetes, a stroke, a heart attack, or to have died. The results for the clinical measurements were mixed. Participants affected by the reform had higher grip strength, lower systolic blood pressure, but they had similar arterial stiffness, height, diastolic blood pressure, intelligence, and happiness to those unaffected by the reform. There was little evidence the reform affected alcohol or tobacco consumption. Participants affected by the reform spent less time watching television, and more time exercising, however, the associations with exercise were imprecise. Finally, participants affected by the reform more likely to earn over £18,000 and £31,000, but were no more likely to earn over £52,000 or £100,000 than those unaffected by the reform. We found some evidence that education may have a larger effect on men’s chances of earning more than £31,000 (p-value for interaction=0.001), but no evidence of interaction with any other outcomes (Tables S1 and S2). There was some evidence that the reform had larger effects on participants predicted to leave before the age of 16: specifically lowering the risk of diabetes, increased likelihood of earning over £31,000, increasing grip strength and happiness, and the likelihood of being diagnosed with a stroke and drank more alcohol (Table S3).

**Table 4:**
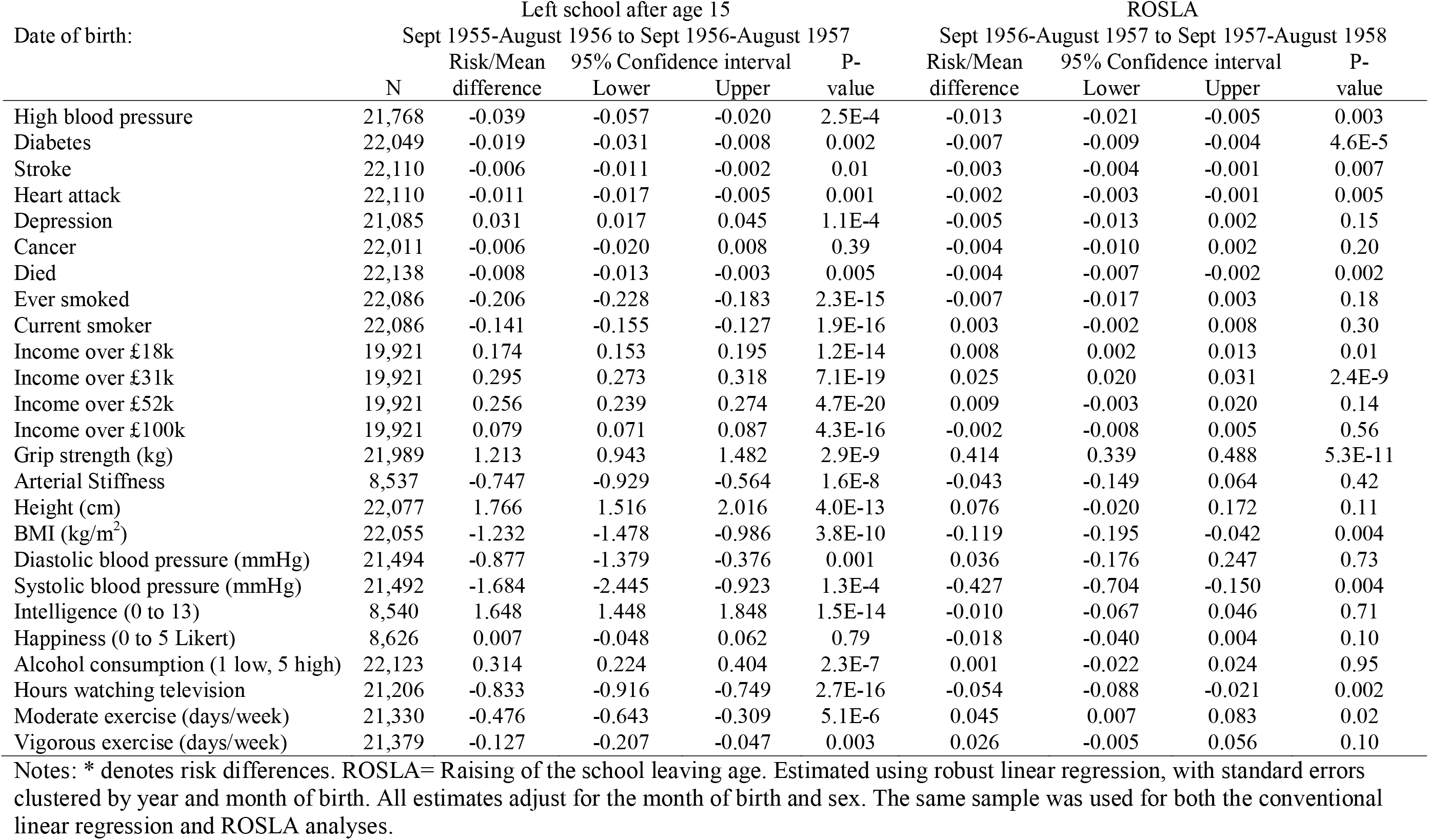
The associations between leaving school after age 15, and attending school after the raising of the school-leaving age (ROSLA) and outcomes.

### Negative control samples

As with the covariate balance tests above we investigated whether the differences in outcomes could be solely explained by the aging process using negative control samples. Participants within each negative control sample experienced the same minimum school leaving ages. Therefore, any differences in these negative control samples are likely to be due to the aging process and not an effect of education. Table 5 reports the placebo results for the negative control samples. These cohorts estimate the average change in each outcome that can be attributed to becoming a year older. On average, the younger participants were less likely to report having high blood pressure, having had a heart attack or depression, had lower systolic and diastolic blood pressure, and arterial stiffness as measured in the clinic. They also reported watching less television. These differences are similar in size to differences in these outcomes observed in Table 4. This suggests these differences are due to the aging process and not an effect of the reform. However, this implies that the aging process cannot entirely explain remaining differences described above, and shown in Table 4 (diagnosed with diabetes or stroke, mortality, grip strength, BMI, and having income over £18,000 or £31,000). The results differed by gender (Tables S4 and S5). For example, there was evidence that on average female participants affected by the reform had 0.15 kg/m^2^ (95%CI: 0.01, 0.30) lower BMI. These gender-specific effects are consistent with the literature.

**Table 5:** Associations of outcomes and the raising of the school-leaving age in two negative control samples.

| Date of birth: | Negative control sample 1 |  |  |  |  | Negative control sample 2 |  |  |  |  |
| --- | --- | --- | --- | --- | --- | --- | --- | --- | --- | --- |
|  | Sept 1955-August 1956 to Sept 1956-August 1957 |  |  |  |  | Sept 1957- August 1958 to Sept 1958-August 1959 |  |  |  |  |
|  | N | Risk/Mean difference | 95% Confidence interval Lower | Upper | P-value | N | Risk/Mean difference | 95% Confidence interval Lower | Upper | P-value |
| High blood pressure | 22,404 | -0.010 | -0.017 | -0.004 | 0.004 | 21,153 | -0.015 | -0.023 | -0.006 | 0.001 |
| Diabetes | 22,685 | -0.001 | -0.003 | 0.001 | 0.49 | 21,376 | -0.002 | -0.005 | 0.001 | 0.25 |
| Stroke | 22,756 | 0.000 | -0.001 | 0.002 | 0.49 | 21,452 | -0.001 | -0.003 | 0.001 | 0.37 |
| Heart attack | 22,756 | -0.002 | -0.004 | 0.000 | 0.04 | 21,452 | 0.002 | 0.000 | 0.004 | 0.04 |
| Depression | 21,658 | 0.012 | 0.004 | 0.019 | 0.004 | 20,428 | -0.004 | -0.009 | 0.002 | 0.18 |
| Cancer | 22,670 | -0.005 | -0.011 | 0.000 | 0.07 | 21,334 | -0.007 | -0.013 | -0.001 | 0.03 |
| Died | 22,789 | 0.002 | -0.001 | 0.004 | 0.16 | 21,476 | 0.001 | 0.000 | 0.002 | 0.14 |
| Ever smoked | 22,728 | -0.011 | -0.023 | 0.001 | 0.07 | 21,436 | -0.001 | -0.012 | 0.010 | 0.88 |
| Current smoker | 22,728 | 0.000 | -0.007 | 0.007 | 0.97 | 21,436 | 0.004 | -0.005 | 0.012 | 0.34 |
| Income over £18k | 20,431 | 0.006 | -0.001 | 0.014 | 0.07 | 19,366 | 0.001 | -0.006 | 0.008 | 0.75 |
| Income over £31k | 20,431 | 0.001 | -0.010 | 0.011 | 0.89 | 19,366 | 0.003 | -0.004 | 0.011 | 0.35 |
| Income over £52k | 20,431 | 0.004 | -0.006 | 0.015 | 0.41 | 19,366 | 0.005 | -0.005 | 0.015 | 0.34 |
| Income over £100k | 20,431 | 0.004 | -0.002 | 0.010 | 0.19 | 19,366 | 0.003 | -0.002 | 0.008 | 0.18 |
| Grip strength (kg) | 22,635 | 0.131 | -0.005 | 0.268 | 0.06 | 21,336 | 0.131 | -0.028 | 0.291 | 0.10 |
| Arterial Stiffness | 8,764 | -0.243 | -0.349 | -0.138 | 8.2E-5 | 8,153 | -0.188 | -0.265 | -0.111 | 4.1E-5 |
| Height (cm) | 22,727 | 0.084 | -0.040 | 0.208 | 0.18 | 21,422 | 0.161 | 0.053 | 0.270 | 0.005 |
| BMI (kg/m<sup>2</sup>) | 22,693 | -0.034 | -0.128 | 0.061 | 0.47 | 21,411 | -0.041 | -0.142 | 0.060 | 0.41 |
| Diastolic blood pressure (mmHg) | 22,101 | -0.288 | -0.517 | -0.059 | 0.02 | 20,829 | -0.310 | -0.525 | -0.095 | 0.007 |
| Systolic blood pressure (mmHg) | 22,099 | -0.904 | -1.242 | -0.567 | 1.2E-5 | 20,829 | -0.636 | -0.864 | -0.409 | 6.8E-6 |
| Intelligence (0 to 13) | 8,755 | -0.076 | -0.146 | -0.006 | 0.03 | 8,163 | -0.063 | -0.129 | 0.002 | 0.06 |
| Happiness (0 to 5 Likert) | 8,847 | 0.008 | -0.016 | 0.032 | 0.50 | 8,227 | -0.008 | -0.028 | 0.012 | 0.44 |
| Alcohol consumption (1 low, 5 high) | 22,778 | -0.012 | -0.034 | 0.010 | 0.28 | 21,461 | -0.030 | -0.054 | -0.005 | 0.02 |
| Hours watching TV | 21,868 | 0.000 | -0.034 | 0.034 | 0.99 | 20,559 | -0.054 | -0.089 | -0.019 | 0.004 |
| Moderate exercise (days/week) | 21,958 | 0.019 | -0.021 | 0.058 | 0.34 | 20,729 | -0.040 | -0.089 | 0.008 | 0.10 |
| Vigorous exercise (days/week) | 21,994 | 0.063 | 0.031 | 0.095 | 5.1E-4 | 20,830 | 0.066 | 0.036 | 0.096 | 1.3E-4 |

Notes: Estimated using robust linear regression, with standard errors clustered by year and month of birth. All estimates adjust for the month of birth and sex.

### Instrumental variables

The associations reported in Table 4 are valid tests of the null hypothesis that education does not affect each outcome. Participants affected by the reform were 14.5 (95%CI: 13.7, 15.2) percentage points more likely to remain in school past age 15 than those who were unaffected. This association is stronger than conventional thresholds for instrumental variable analysis (F-statistic=1393). In Table 6 we report instrumental variable estimates of the size of the effects of remaining in school past the age of 15. These results suggest that remaining in school reduced the probability of being diagnosed with diabetes, stroke, or dying by 4.5 (95%CI: 2.7 to 6.3), 1.7 (95%CI: 0.6 to 2.8), and 2.9 (95%CI: 1.3 to 4.5) percentage points. These estimates are larger than the conventional linear regression (Hausman test for differences p=9.7E-7, p=0.002, and p=3.4E-4). Staying in school also increased the participants grip strength by 2.89 (95%CI: 2.38 to 3.39) kg, which again was more than implied by the conventional linear regression (Hausman p-value=3.6E-29). Finally, the instrumental variable results imply that staying in school increases the likelihood of earning more than £18,000 or £31,000 by 5.6 (95%CI: 1.5 to 9.6) and 18.8 (95%CI: 14.9 to 22.6) percentage points. Both of these estimates are smaller than implied by conventional linear regression (Hausman p-values=0.001). These results exceeded the Benjamini and Hochberg (1995) false discovery rate threshold at δ=0.05 across 25 outcomes.(28) Figures 5 and 6 show the differences between the conventional linear regression results and the instrumental variable results. They plot the point estimate and confidence intervals for the conventional linear regression and the instrumental variable results. Tables S6 and S7 report the instrumental variable results stratified by gender.

**Table 6:** The effects of leaving school after age 15, instrumental variable regression (left) and conventional regression (right)

|  | Instrumental variable regression |  |  |  |  |  | Conventional linear regression |  |  |  |
| --- | --- | --- | --- | --- | --- | --- | --- | --- | --- | --- |
|  | N | Risk/Mean difference | 95% Confidence interval |  | P-value | Hausman p-value | Partial F-stat | Risk/Mean difference | 95% Confidence interval |  |
|  |  |  | Lower | Upper |  |  |  |  | Lower | Upper |
| High blood pressure | 21,768 | -0.091 | -0.143 | -0.040 | 5.5E-4* | 0.08 | 1327 | -0.039 | -0.057 | -0.020 |
| Diabetes | 22,049 | -0.045 | -0.063 | -0.027 | 9.7E-7* | 0.02 | 1318 | -0.019 | -0.031 | -0.008 |
| Stroke | 22,110 | -0.017 | -0.028 | -0.006 | 0.002* | 0.10 | 1327 | -0.006 | -0.011 | -0.002 |
| Heart attack | 22,110 | -0.012 | -0.019 | -0.004 | 0.001* | 0.87 | 1327 | -0.011 | -0.017 | -0.005 |
| Depression | 21,085 | -0.038 | -0.087 | 0.012 | 0.13 | 0.01 | 1221 | 0.031 | 0.017 | 0.045 |
| Cancer | 22,011 | -0.028 | -0.068 | 0.012 | 0.17 | 0.30 | 1315 | -0.006 | -0.020 | 0.008 |
| Died | 22,138 | -0.029 | -0.045 | -0.013 | 3.4E-4* | 0.009 | 1330 | -0.008 | -0.013 | -0.003 |
| Ever smoked | 22,086 | -0.048 | -0.116 | 0.019 | 0.16 | 0.001 | 1327 | -0.206 | -0.228 | -0.183 |
| Current smoker | 22,086 | 0.018 | -0.015 | 0.050 | 0.28 | 2.0E-5 | 1327 | -0.141 | -0.155 | -0.127 |
| Income over £18k | 19,921 | 0.056 | 0.015 | 0.096 | 0.007* | 0.001 | 1112 | 0.174 | 0.154 | 0.194 |
| Income over £31k | 19,921 | 0.188 | 0.149 | 0.226 | 1.7E-21* | 0.001 | 1112 | 0.295 | 0.267 | 0.323 |
| Income over £52k | 19,921 | 0.063 | -0.015 | 0.141 | 0.11 | 6.3E-4 | 1112 | 0.256 | 0.237 | 0.275 |
| Income over £100k | 19,921 | -0.014 | -0.058 | 0.030 | 0.54 | 0.002 | 1112 | 0.079 | 0.070 | 0.088 |
| Grip strength (kg) | 21,989 | 2.885 | 2.381 | 3.390 | 3.6E-29* | 3.1E-4 | 1302 | 1.213 | 0.943 | 1.482 |
| Arterial Stiffness | 8,537 | -0.323 | -1.067 | 0.422 | 0.40 | 0.27 | 469 | -0.747 | -0.929 | -0.564 |
| Height (cm) | 22,077 | 0.527 | -0.088 | 1.143 | 0.09 | 0.002 | 1324 | 1.766 | 1.516 | 2.016 |
| BMI (kg/m<sup>2</sup>) | 22,055 | -0.820 | -1.323 | -0.316 | 0.001* | 0.18 | 1324 | -1.232 | -1.478 | -0.986 |
| Diastolic blood pressure (mmHg) | 21,494 | 0.249 | -1.110 | 1.607 | 0.72 | 0.10 | 1275 | -0.877 | -1.379 | -0.376 |
| Systolic blood pressure (mmHg) | 21,492 | -2.975 | -4.740 | -1.209 | 9.6E-4* | 0.11 | 1274 | -1.684 | -2.445 | -0.923 |
| Intelligence (0 to 13) | 8,540 | -0.078 | -0.482 | 0.326 | 0.71 | 1.1E-5 | 469 | 1.648 | 1.448 | 1.848 |
| Happiness (0 to 5 Likert) | 8,626 | -0.139 | -0.290 | 0.012 | 0.07 | 0.08 | 480 | 0.007 | -0.048 | 0.062 |
| Alcohol consumption (1 low, 5 high) | 22,123 | 0.005 | -0.143 | 0.153 | 0.95 | 0.002 | 1329 | 0.314 | 0.224 | 0.404 |
| Hours watching TV | 21,206 | -0.372 | -0.577 | -0.167 | 3.8E-4* | 0.001 | 1292 | -0.833 | -0.916 | -0.749 |
| Moderate exercise (days/week) | 21,330 | 0.327 | 0.073 | 0.581 | 0.01* | 3.0E-4 | 1211 | -0.476 | -0.643 | -0.309 |
| Vigorous exercise (days/week) | 21,379 | 0.186 | -0.022 | 0.395 | 0.08 | 0.02 | 1205 | -0.127 | -0.207 | -0.047 |

Notes: Estimated using robust linear regression, with standard errors clustered by year and month of birth. All estimates adjust for the month of birth and sex. \* Exceeds Benjamini and Hochberg (1995) corrected threshold for false discovery rate at $\delta=0.05$ across 25 outcomes.(28)

**Figure 5:**
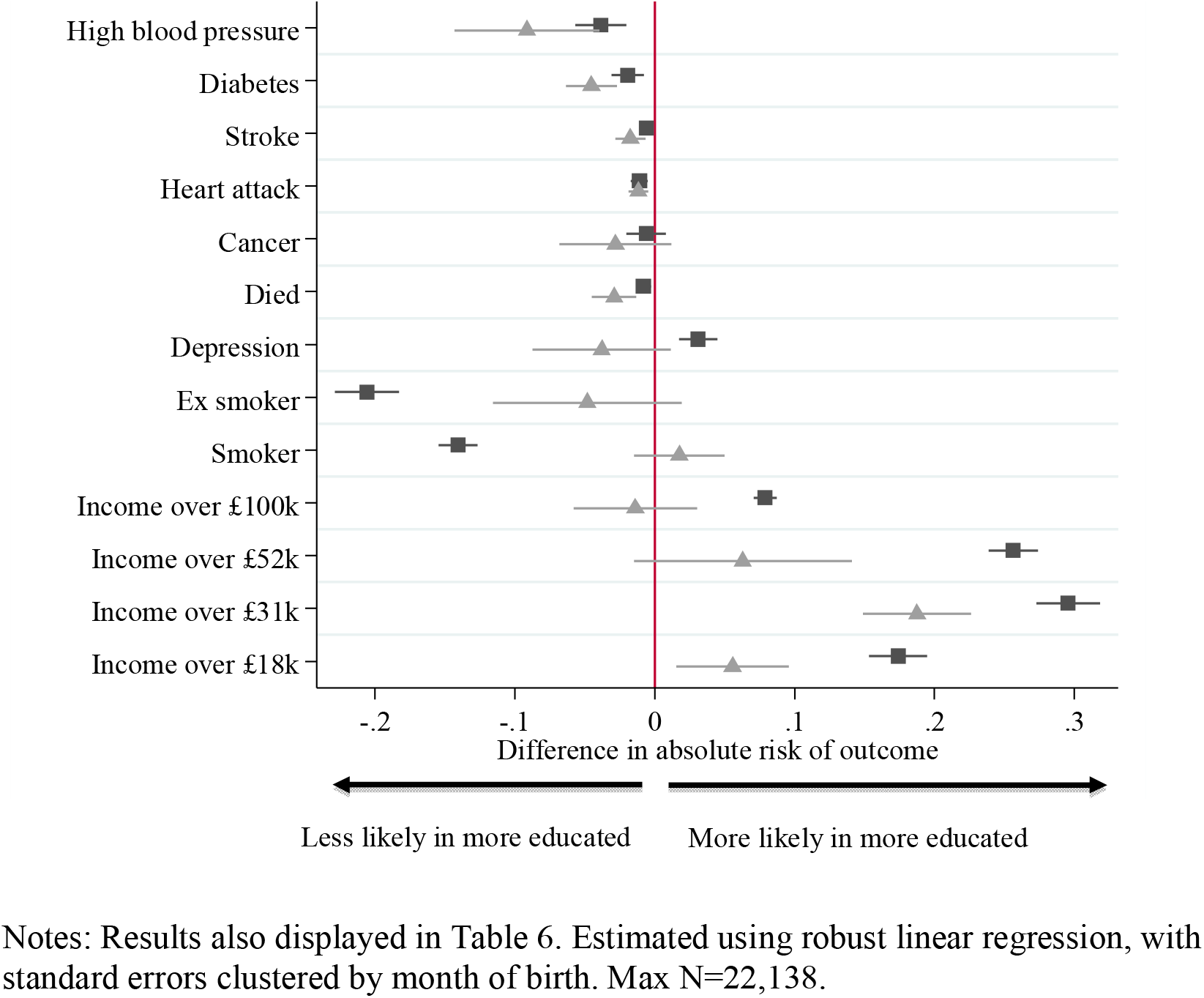
Difference in risk of outcomes between those who left school after age 15. Estimated by actual education attainment (squares), and using the raising of the school-leaving age as an instrument (triangles).

**Figure 6:**
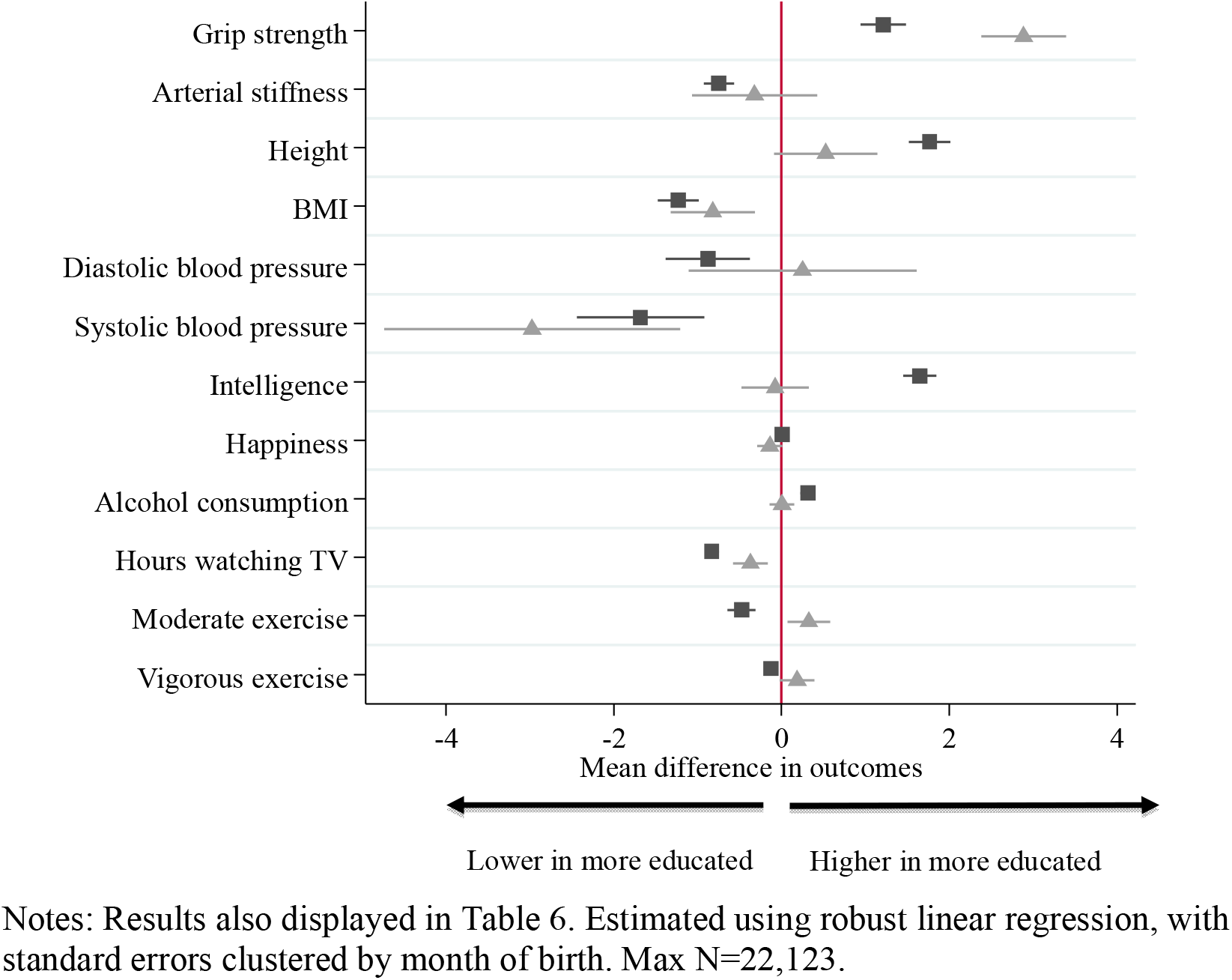
Difference in mean outcomes between those who left school after age 15. Estimated by actual education attainment (squares), and using the raising of the school-leaving age as an instrument (triangles).

## Discussion

This study provides some of the strongest evidence to date about the causal effects of education. First, our instrumental variable results indicate that socioeconomic and genomic factors can explain some of the observed associations of educational attainment and outcomes later in life. In contrast to the observational associations, we found little evidence that education had causal effects on smoking, alcohol consumption, arterial stiffness, intelligence, or likelihood of having an annual income over £100,000. Second, our results provide robust evidence that education is likely to have a causal effect on other outcomes, including diagnoses of diabetes, stroke, mortality, grip strength and having incomes above £18,000 and £31,000. Finally, we found molecular genetic evidence that traditional approaches to investigate the effects of educational attainment, such as multivariable adjusted regression, are likely to suffer from residual genomic confounding.

Clark and Royer found the participants of the Health Survey for England and the General Household Survey affected by the reform were by 26.1 (95%CI: 23.0 to 29.2) percentage points more likely to stay in school after age 15.(1) We found a smaller difference (14.0, 95%CI: 12.9 to 15.1), this may be because on average the participants of UK Biobank are more educated. Clark and Royer found little evidence that the raising of the school-leaving age reduced mortality (odds-ratio=0.99, 95%CI: 0.94 to 1.05). Whereas we found people affected by the reform had a lower risk of death (odds-ratio=0.64, 95%CI: 0.50 to 0.83). We found little evidence of reductions in mortality in the two negative control samples odds-ratios= 1.22 (95%CI: 0.97 to 1.53) and 1.08 (95%CI: 0.91 to 1.29) for negative control samples 1 and 2, in the two years before and the two years after the reform. Clark and Royer found little evidence of effects of the reform on self-reported health. This may be because they used relatively liberal bandwidths, up to 138 months either side of the reform. This meant they were comparing individuals born up to 11.5 years before the reform, to individuals who were born up to 11.5 years after the reform. For some, but not all, analyses they allowed for a month and year of birth trend, which was allowed to change after the reform. The advantage of using such a liberal bandwidth is that they could include more data, and to obtain sufficiently precise results. However, this is potentially at the expense of introducing bias by comparing individuals who differ because of secular trends. We obtained similar precision to Clark and Royer using a highly conservative 12-month bandwidth. As a sensitivity analysis, we provide results using a comparable specification to Clark and Royer in the supplementary materials, see Tables S8, S9, and S10. Furthermore, the majority of their health outcomes were self-reported in a general health survey, whereas we had precise clinic measures of specific measures of gaining which are known to change with age, such as grip strength. This means our outcomes are likely to suffer from less measurement error. Our data were consistently measured in a single study across a relatively narrow time window, whereas previous results pooled data from many years. Again this means our results are likely to suffer from less measurement error.

Epidemiologists have argued that education has causal effects on intelligence later in life. For example, Richards and Sacker found that educational attainment by age 26 was associated with intelligence at age 53,(29) which they argue was evidence that education had a causal effect on intelligence.(16) However, Deary and Johnson raised doubts about this interpretation and called for greater clarity about the assumptions underlying these analyses.(19) We found little evidence of a causal effect of education on intelligence later in life. Nguyen and colleagues used increases in the legal school-leaving ages in the United States to investigate the effects of education on risk of dementia later in life.(21) They found evidence that education reduced the risk of dementia. We cannot test this hypothesis in the UK Biobank because too few participants have been diagnosed with dementia.

People with more education are much less likely to smoke. However, it is not clear whether this is due to a causal effect of education. Gilman and colleagues found the association between education and smoking status was attenuated in sibling fixed effects designs.(30) We found that participants who remained in school were 20.1 percentage points less likely to have ever smoked. However, we found little evidence that the raising of the school leaving age affected smoking behavior. We also found that educated participants drank more heavily, but there was little evidence that this was caused by education. This is consistent with recent evidence from other UK surveys. Silles (2015) used data from the General Household Survey for Great Britain and found that whilst people who left school at an older age were less likely to smoke, there were few differences in ever-smoking rates among individuals unaffected and affected by the 1947 and 1972 raising of the school leaving age.(31) We also found some evidence that the effects of the reform were greatest in participants who would otherwise have been expected to leave at age 15.

### Strengths and limitations

A key strength of our study is that we used a natural experiment to identify the effects of education. The raising of the school-leaving age in 1972 provided exogenous variation in the length of schooling people received. We found few pre-existing differences between participants on either side of the reform, suggesting that it can be used as a potentially valid instrumental variable.(32) A strength of our study is it uses one of the largest samples to date to investigate the effects of education on a wide range of outcomes. Our outcomes were recorded both in interviews and via linked Office of National Statistics registry data. This means our outcomes are likely to suffer from relatively little measurement error. Furthermore, we were able to restrict our sample to people born in England who were affected by the reform. In addition, we used genome-wide data to show that conventional multivariable adjusted regression results are likely to suffer from genomic confounding. A potential limitation of our study is that our treatment group, people affected by the reform, are one year younger than our control group, those unaffected by the reform. Many of the outcomes we investigated increase linearly or log-linearly over time. This means it is difficult to determine if any differences we observed are due to an additional year of aging or the reform. We addressed this by using negative control samples to estimate the average effects of aging. For the results for outcomes such as blood pressure, where we observed similar differences in the negative control samples as the main results, are likely to be due to aging and not an effect of the reform. However, it is likely the reform affected outcomes, such as income, where we observed much larger differences in the main results than in the negative control samples. Further, aging cannot explain why we found little evidence of causal effects of staying in school on intelligence, smoking, and alcohol consumption.

Whilst a representative sample is not a necessary condition for making causal inferences,(33) collider bias could affect our results because Biobank is a volunteer sample, which over-represents highly educated people. People affected by the reform may be more likely to participate in the study.(34) Therefore less educated individuals, who would have stayed in school had they attended school after the reform (the compliers), may be under-represented in the pre-reform sample. If true, this could attenuate our results towards the null, because these marginal students would reduce the average outcome in the “treatment” group, and be missing from the “control” group. This would improve the control group’s outcome relative to the treatment group.

Nevertheless, our results are consistent with previous results which used random samples.(31) So collider bias is unlikely to explain why there was little evidence that the reform affected intelligence, smoking and alcohol consumption. However, this issue warrants further investigation in future research. There was limited time to collect measures during the participants’ assessment center visits, therefore our measure of intelligence is relatively coarse. However, despite this, participants who remained in school had substantially higher intelligence. Our instrumental variable estimates of the effects of schooling on intelligence are sufficiently precise to rule out even relatively small effects on intelligence. Finally, our instrumental variable results are estimates of the local average treatment effect of schooling.(35) Specifically, they are the estimates of the causal effects of being forced to remain in school after the age of 15, on those who would otherwise have left school. These effects may not be externally valid to infer either the effects of compelling students to remain in school for longer or of the effects on other populations.(36, 37) This means these results may not be valid estimates of the effect of education on “always takers”, that is people who would always remain in school regardless of the reform. Nevertheless, our results are likely to be internally valid estimates of the effects of schooling on people affected by the reform.

## Conclusions

Does education have causal effects? Yes, whilst education is not the panacea implied by naïve multivariable adjusted regression, in this sample staying in school did result in substantial benefits. We found robust evidence that staying in school is likely to have causal effects on some, but not all health and socioeconomic outcomes later in life. This adds to our understanding of the long-term consequences of educational decisions in childhood and adolescence. We resolve two debates about the causal effects of schooling. First, our results suggest education is unlikely to cause differences in intelligence later in life.(16, 19) Second, previous research using these reforms found little evidence that education affected health, may be due to poor measurement of health outcomes and inadequate statistical power.(1)

## Materials and Methods

### Data

We used data from 502,624 participants of the UK Biobank project.(25) The participants, aged between 40 and 69, were originally recruited between 2006 and 2010. In our primary analysis, we restricted our sample to participants were born in England in the school cohorts in years immediately before and after the reform took place. We do this because we have a large enough sample born in these years to precisely identify the effects of schooling.

### Exposure: left school after age 15

The participants were asked if they had a college or university degree. If they did not have a degree they were asked what age they left full-time education. We coded participants who reported having a degree as leaving full-time education at age 21.

## Outcomes

### Health outcomes

The participants were asked whether they had ever been diagnosed by a doctor with the following health conditions: high blood pressure, stroke, diabetes, or heart attack. They were asked if they had ever had a whole week where they felt depressed or down. The death of the participants was defined using linked Office of National Statistics mortality data. Follow-up for the linked mortality data started with the first death on 10^th^ May 2006 ended with the last recorded death on 17^th^ February 2014. The diagnoses of cancer and information were taken from the cancer registry. The first recorded cancer diagnosis was on 20^th^ September 1957 and the last on 25^th^ October 2013.

### Height, BMI, Blood pressure, atrial stiffness, grip strength, and intelligence

Height and weight were measured during the participants’ visit to a UK Biobank assessment center. Two measures of diastolic and systolic blood pressure were recorded via an electronic blood pressure monitor. The measurements were taken two minutes apart. Atrial stiffness was measured using an electronic measure device. Grip strength was measured in kilos using a hydraulic hand dynamometer. We residualized the measures of grip strength and atrial stiffness to control for potential between device heterogeneity. Fluid intelligence was measured via 13 logic puzzles that the participants had to answer in 2 minutes. Their score is the number of correct answers.

### Health behaviors and income

During their assessment center visit, the participants were asked to report their health behaviors. They were asked about how frequently they consumed alcohol. This is coded 6 if they drank every day, 5 for three or four times a week, 4 for once or twice a week, 3 for one to three times a week, 2 for special occasions only, and 1 for never. They were asked if they smoked, or had ever smoked. They were asked how often they vigorously and moderately exercised in a typical week. Finally, they were asked if their pre-tax income was below £18,000; between £18,000 and £30,999; between £31,000 and £50,999; between £52,000 and £100,000; or above £100,000. Participants who did not answer these questions were coded as missing.

### Genotype data

The participants provided a blood sample. This sample was used to extract DNA and genotype using the Axiom and BiLEVE genome-wide arrays. These arrays genotyped around 800,000 single nucleotide polymorphism (SNPs) for each participant. The genotyping data was used to impute SNPs which were not directly genotyped using the 1000 genomes and UK10K reference panels. The imputation produced a likelihood of each participant having a specific genotype at each (e.g. AA=0.1, TA=0.9, and TT=0). This resulted in a dataset of around 80,000,000 SNPs. For each participant, we created a genome-wide allele score by summing the number of genetic variants they had that were associated with higher educational attainment. We weighted each variant by its association with education reported in a large genome-wide association study.(27) This study reported the association of 8,259,394 genetic variants and the years of education in a meta-analysis of 64 studies, which did not include the UK Biobank. We normalized the allele score have mean zero and standard deviation one. The allele score represents the effects of the known effects of genetic variants on educational attainment. However, these scores only explain a minority (r^2^=1.32% in the full Biobank sample) of the variation in educational attainment explained by genome-wide data.(27, 38, 39) This is because of limited statistical power of existing genome-wide association studies of educational attainment. One consequence of this is that the genetic score is too poor a proxy for the total genetic effects on educational attainment to be used as a conventional covariate in a regression. However, this score does provide a valid test of the null-hypothesis that previously unobserved genetic covariates do not confound the association of education and other outcomes.(32)

### Statistical methods

We use the changes in the school-leaving age to identify the effects of schooling on a range of outcomes. Our empirical strategy has five steps. First, we estimated the effect of the reforms on the proportion of participants who remained in school after age 15. Second, we investigated the associations of potential confounders with educational attainment and across the cohorts affected by the reform.(32) Third, we estimated the reduced form associations of the reform and the outcomes. Fourth, we used instrumental variable estimators to estimate the effects of the attending school on each of the outcomes. For continuous outcomes, we used conventional two-stage least squares,(40) for binary outcomes we used semi-parametric additive structural mean models.(41) To address concerns about multiple hypothesis testing, we report whether the instrumental variable results for each outcome exceed a Benjamini and Hochberg (1995) false discovery rate threshold at δ=0.05 across 25 outcomes.(28) Fifth, we conducted two negative control analyses.(28) We did this by constructing two cohorts of students, born in the two school years before and the two years after the reform. Analogous with the primary cohort we designated participants born in the more recent of the two school years as being exposed to the “placebo” reform. There are no institutional differences between each year, therefore any differences observed in these negative control cohorts will be due age effects and not an effect of raising the school-leaving age.

### Identification

The raising of the school-leaving age will be a valid natural experiment for testing whether remaining in school at age 15 affects later outcomes under the following three assumptions. First, participants who attended school after the leaving age was increased must be more likely to stay in school. Second, there must be no pre-existing differences between the cohort who attended school in the year immediately before and immediately after the reform. Finally, the reform must not have any other direct effects on the outcomes. We can test the first assumption by testing whether participants affected by the reform are more likely to stay in school. We can falsify the second assumption by investigating if there were any pre-existing differences between those affected and unaffected by the reform. The final assumption cannot be empirically tested, and could be invalid if the reform also affected the labor market around the time that the participants entered the workforce. However, claimant count statistics for the UK show that the cohorts entering the labor force immediately before and after the reform faced broadly similar conditions, with increases in unemployment related to the oil crises of the 1970s not being seen until 1975 onwards.(42, 43)

#### 1. The effect of increasing the minimum leaving age on educational attainment

We used a regression discontinuity design to estimate the effects of increasing the school-leaving age from age 15 to 16 on the proportion of students who report leaving school before the age of 15. To investigate the effect of the reform on school attendance we estimated a regression of staying school after age 15 on a dummy variable equal to one if the participant was a member of the cohort affected by the reform, and equal to zero if they were not affected. In this and all subsequent analyses we included covariates for the month of birth, to control for seasonality, and gender. In contrast to Clark and Royer (2013), we do not include a term for birth cohort because we restricted our sample to people born in the single school years immediately before and after the reform. The regression discontinuity design is identified by assuming that the reform is independent of the unobserved confounding factors, and has no other direct effects on the outcome. The effect of the reform on the probability of participants staying in school after the age of 15, our parameter of interest, is the effect of remaining in school on those who were affected by the reform. This is a local average treatment effect.(35) We report this parameter on the risk difference scale. Our regressions allow for general form heteroskedasticity and clustering by year and month of birth.

#### 2. Covariate balance tests

We compared the associations of eleven potential confounders *C*_*ict*_ and the exposure, left school after the age of 15, *E*_*ict*_, and the indicator of the reform, *D*_*ic*_. We estimated these associations conditional on the same set of covariates, 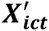, as above and the standard errors allow for clustering by year and month of birth.

#### 3. Effects of increasing the school-leaving age on outcomes in later life

##### A. Reduced form

We estimated the associations of leaving school after age 15 and the outcomes and the association of the reform and each of the outcomes using the following linear regressions:

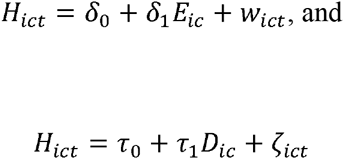

The first is a linear regression of each of the health outcomes on whether the participant stayed in school after the age of 15. The second regression is the reduced form association of the health outcomes and the reform. As above, each regression includes terms for gender and month of birth to account for the season of birth. The reduced form is a valid test of the null-hypothesis that schooling does not affect the outcomes.

We tested whether the reform had larger effects on people who would otherwise have been expected to leave school at age 15. We estimated the probability that a participant would remain in school after the age of 15 using logistic regression and data from individuals born before 31^st^ August 1956. This model included indicators for the participants’ assessment center, year and month of birth, sex, whether mother smoked during pregnancy, were breastfed, their relative height and body size at age 8, number of brothers and sisters, the normalized genome-wide education score, and their ethnicity. Missing data were replaced at the mean and indicators variables for missing values were included. We estimated the following regression:

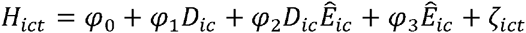

Where 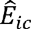 is probability of remaining in education from the logistic regression. For each outcome we report the coefficients on the reform indicator, and the coefficient on the interaction term and the effect of the reform. The effect of the reform on participants predicted to leave is indicated by *φ*_1_, and the effect on those expected to stay is indicated by *φ*_1_ + *φ*_3_. As with the main results above we adjust for sex and month of birth, and the interaction of these variables with predicted education. (44)

##### B. Instrumental variables

We estimated the causal effect of schooling using instrumental variables estimators. For the continuous outcomes, we estimated mean differences using Two-Stage Least Squares (2SLS),(40) and for the binary outcomes we estimated risk differences using additive structural mean models.(41) These models can be identified by making one of three assumptions.(41) First, for the continuous outcomes we could assume that staying in school has the same effect on the outcomes for all participants. This identifies the average effects of staying in school but is implausible for binary outcomes.(45) Second, for the binary outcomes, we could assume a monotonic relationship between the reform and the participants’ likelihood of staying in school after the age of 15. In the potential outcomes framework, we assumed that *E*[*Y*(1) – *Y*(0) | *E*(1) – *E*(0) > 0]. This requires that there were no participants who were “defiers”, who would have stayed in school if they were not affected by the reform, but would have left school if they were affected by the reform. Under monotonicity, the instrumental variable estimators estimate a local average treatment effect. This is the effects of treatment in the subgroup of participants whose decisions were affected by the reform.(40) That is the people in the year after the reform who would have chosen to leave school at 15 had the reform not been introduced. Finally, we could assume that the effects of education are not affected by the reform (no effect modification). This would identify the effects of education on participants who chose to stay in school. We report the partial F-statistic of the association of remained in school *E*_*ict*_ and the reform *D*_*ic*_. We also report the test for endogeneity (using a C-statistic, which is a heteroskedasticity robust Hausman test (46, 47), that *E*[*E*_*ict*_*w*_*ict*_] = 0. This implicitly tests for differences between the linear regression and instrumental variable estimates.(47) All estimates allow for clustered standard errors by year and month of birth and include controls for gender.

##### C. Regression discontinuity design

Finally, we used a regression discontinuity design with variable bandwidths to investigate the robustness of our findings. This is a fuzzy regression discontinuity design, as the reform only increased the probability of staying in school.(48) In our main analysis above we present reduced forms on the participants born between September 1956 and August 1958. This is a regression discontinuity design estimates mean differences with a bandwidth of one year. We used the rd command in Stata to produce local linear regression discontinuity estimates for larger bandwidths as a sensitivity analysis.(49)

##### D. Negative control samples

We were concerned that differences between the two school years may occur because of the participants affected by the reform were a year younger on average than participants unaffected by the reform. To investigate this, we investigated the effects of two placebo reforms. The first placebo reform compares the cohort which left school two years before the reform was introduced (i.e. those born between September 1955 and August 1956) to the cohort that left school in the year before the reform was introduced (i.e. those born between September 1956 to August 1957). The second compares participants in the school cohort in the year after the reform, (i.e. those born between September 1957 to August 1958) to those in the cohort two years after the reform (i.e. those born between September 1958 to August 1959). These cohorts have the same average difference in age as the two school years either side of the reform, but they experienced the same school-leaving age. Therefore, any differences between the younger and older school years in these negative control samples are likely to be due to aging and will not be due to the effects of education. We estimated the reduced form associations using these negative control samples. We repeated all of the analyses stratified by sex as a sensitivity analysis and report these results in the online appendix.

All analyses were conducted in StataMP 14.0.(50) Code used to generate these results can be found at (https://github.com/nmdavies/UKbiobankROSLA) and the data used has been archived with UK Biobank (http://www.ukbiobank.ac.uk) and can be accessed by contacting the study. The protocol for this study is available as an online appendix.

## Acknowledgements

We would like to thank the Social Science Genetic Association Consortium for providing the coefficients from the Educational attainment GWAS, and Gibran Hemani, Lavinia Paternoster, David Carslake, Jack Bowden, Louisa Zuccolo Evie Stergiakouli and Eleanor Sanderson for helpful comments on an earlier draft. All mistake remain our own.

---

**Table S1:**
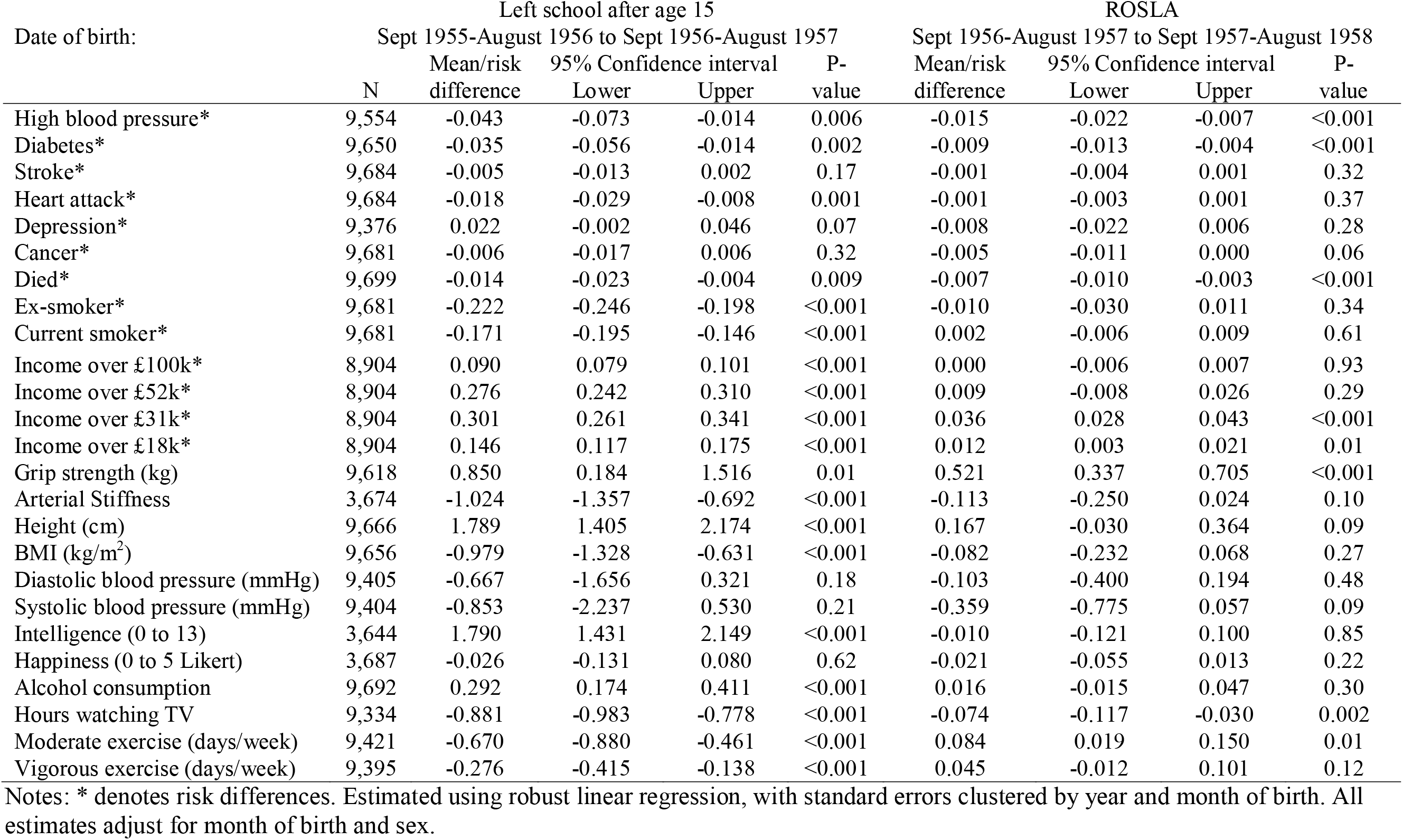
The associations between leaving school after age 15, and attending school after the raising of the school leaving age (ROSLA) and outcomes <u>for MALES</u>.

**Table S2:** The associations between leaving school after age 15, and attending school after the raising of the school leaving age (ROSLA) and outcomes <u>for FEMALES</u>.

| Date of birth: | Left school after age 15 |  |  |  |  | ROSLA |  |  |  |
| --- | --- | --- | --- | --- | --- | --- | --- | --- | --- |
|  | Sept 1955-August 1956 to Sept 1956-August 1957 |  |  |  |  | Sept 1956-August 1957 to Sept 1957-August 1958 |  |  |  |
|  | N | Mean/risk difference | 95% Confidence interval<br>Lower | Upper | P-value | Mean/risk difference | 95% Confidence interval<br>Lower | Upper | P-value |
| High blood pressure* | 12,214 | -0.035 | -0.054 | -0.015 | 0.001 | -0.012 | -0.025 | 0.002 | 0.09 |
| Diabetes* | 12,399 | -0.007 | -0.015 | 0.001 | 0.08 | -0.005 | -0.008 | -0.002 | 0.003 |
| Stroke* | 12,426 | -0.007 | -0.012 | -0.001 | 0.02 | -0.003 | -0.005 | -0.002 | <0.001 |
| Heart attack* | 12,426 | -0.005 | -0.009 | 0.000 | 0.03 | -0.002 | -0.004 | -0.001 | <0.001 |
| Depression* | 11,709 | 0.038 | 0.022 | 0.054 | <0.001 | -0.004 | -0.015 | 0.007 | 0.50 |
| Cancer* | 12,330 | -0.006 | -0.025 | 0.012 | 0.50 | -0.003 | -0.011 | 0.005 | 0.43 |
| Died* | 12,439 | -0.004 | -0.008 | 0.001 | 0.12 | -0.002 | -0.005 | 0.001 | 0.11 |
| Ex-smoker* | 12,405 | -0.192 | -0.227 | -0.157 | <0.001 | -0.004 | -0.013 | 0.004 | 0.30 |
| Current smoker* | 12,405 | -0.116 | -0.134 | -0.099 | <0.001 | 0.003 | -0.003 | 0.010 | 0.29 |
| Income over £100k* | 11,017 | 0.070 | 0.059 | 0.081 | <0.001 | -0.003 | -0.014 | 0.007 | 0.53 |
| Income over £52k* | 11,017 | 0.240 | 0.222 | 0.258 | <0.001 | 0.009 | -0.002 | 0.021 | 0.12 |
| Income over £31k* | 11,017 | 0.290 | 0.255 | 0.326 | <0.001 | 0.018 | 0.011 | 0.025 | <0.001 |
| Income over £18k* | 11,017 | 0.196 | 0.164 | 0.228 | <0.001 | 0.004 | -0.004 | 0.012 | 0.32 |
| Grip strength (kg) | 12,371 | 1.501 | 1.167 | 1.836 | <0.001 | 0.336 | 0.189 | 0.482 | <0.001 |
| Arterial Stiffness | 4,863 | -0.550 | -0.759 | -0.341 | <0.001 | 0.000 | -0.127 | 0.127 | 1.00 |
| Height (cm) | 12,411 | 1.756 | 1.403 | 2.110 | <0.001 | 0.008 | -0.091 | 0.106 | 0.87 |
| BMI (kg/m<sup>2</sup>) | 12,399 | -1.425 | -1.761 | -1.088 | <0.001 | -0.146 | -0.304 | 0.011 | 0.07 |
| Diastolic blood pressure (mmHg) | 12,089 | -1.054 | -1.540 | -0.568 | <0.001 | 0.140 | -0.211 | 0.492 | 0.42 |
| Systolic blood pressure (mmHg) | 12,088 | -2.345 | -3.267 | -1.424 | <0.001 | -0.480 | -0.964 | 0.003 | 0.05 |
| Intelligence (0 to 13) | 4,896 | 1.546 | 1.351 | 1.740 | <0.001 | -0.011 | -0.091 | 0.070 | 0.79 |
| Happiness (0 to 5 Likert) | 4,939 | 0.031 | -0.034 | 0.097 | 0.33 | -0.016 | -0.042 | 0.010 | 0.21 |
| Alcohol consumption | 12,431 | 0.328 | 0.240 | 0.416 | <0.001 | -0.011 | -0.057 | 0.035 | 0.62 |
| Hours watching TV | 11,872 | -0.796 | -0.912 | -0.679 | <0.001 | -0.041 | -0.081 | 0.000 | 0.05 |
| Moderate exercise (days/wk) | 11,909 | -0.317 | -0.493 | -0.141 | 0.001 | 0.014 | -0.056 | 0.083 | 0.69 |
| Vigorous exercise (days/wk) | 11,984 | -0.007 | -0.131 | 0.116 | 0.90 | 0.009 | -0.033 | 0.051 | 0.67 |

Notes: \* denotes risk differences. Estimated using robust linear regression, with standard errors clustered by year and month of birth. All estimates adjust for month of birth and sex.

**Table S3:**
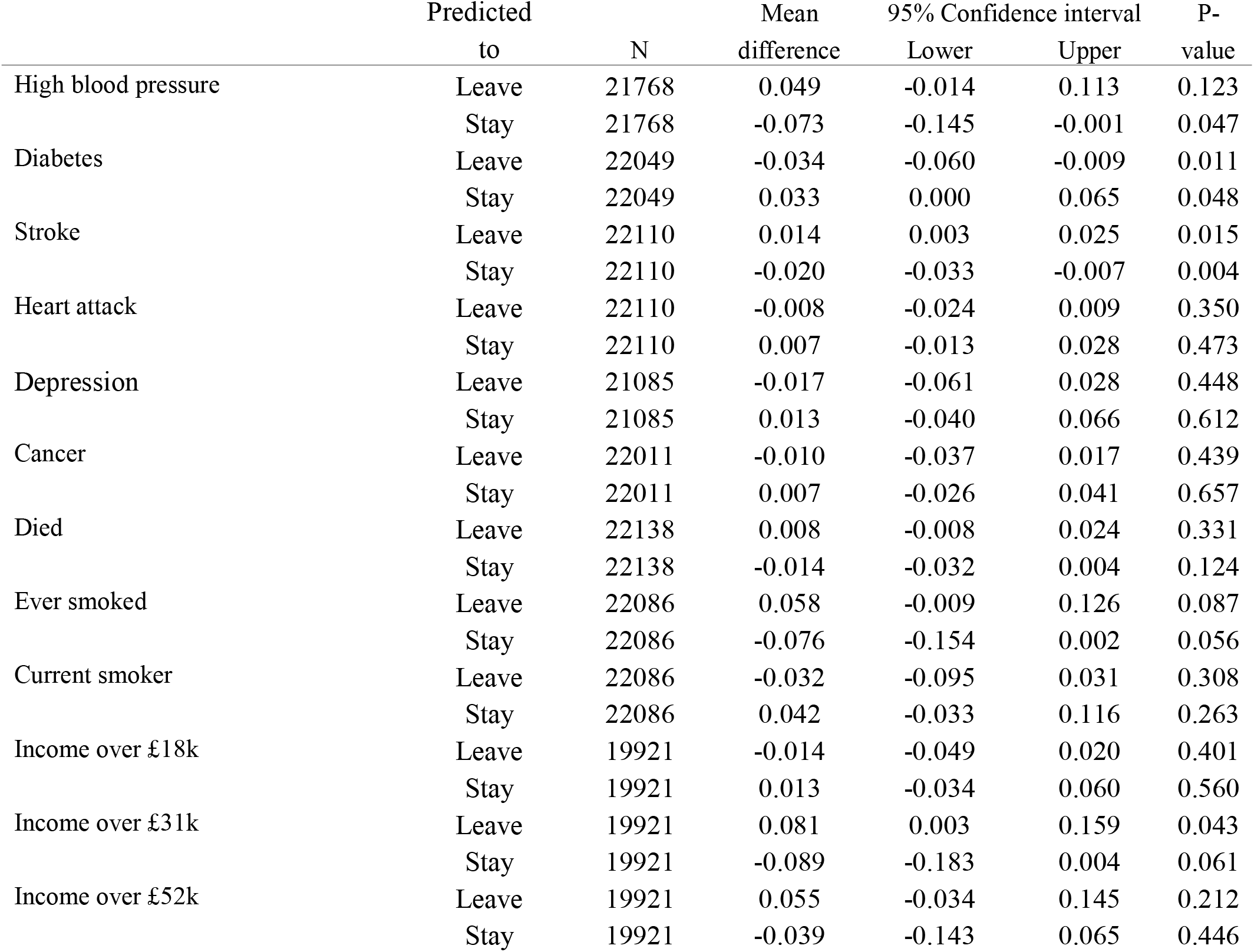
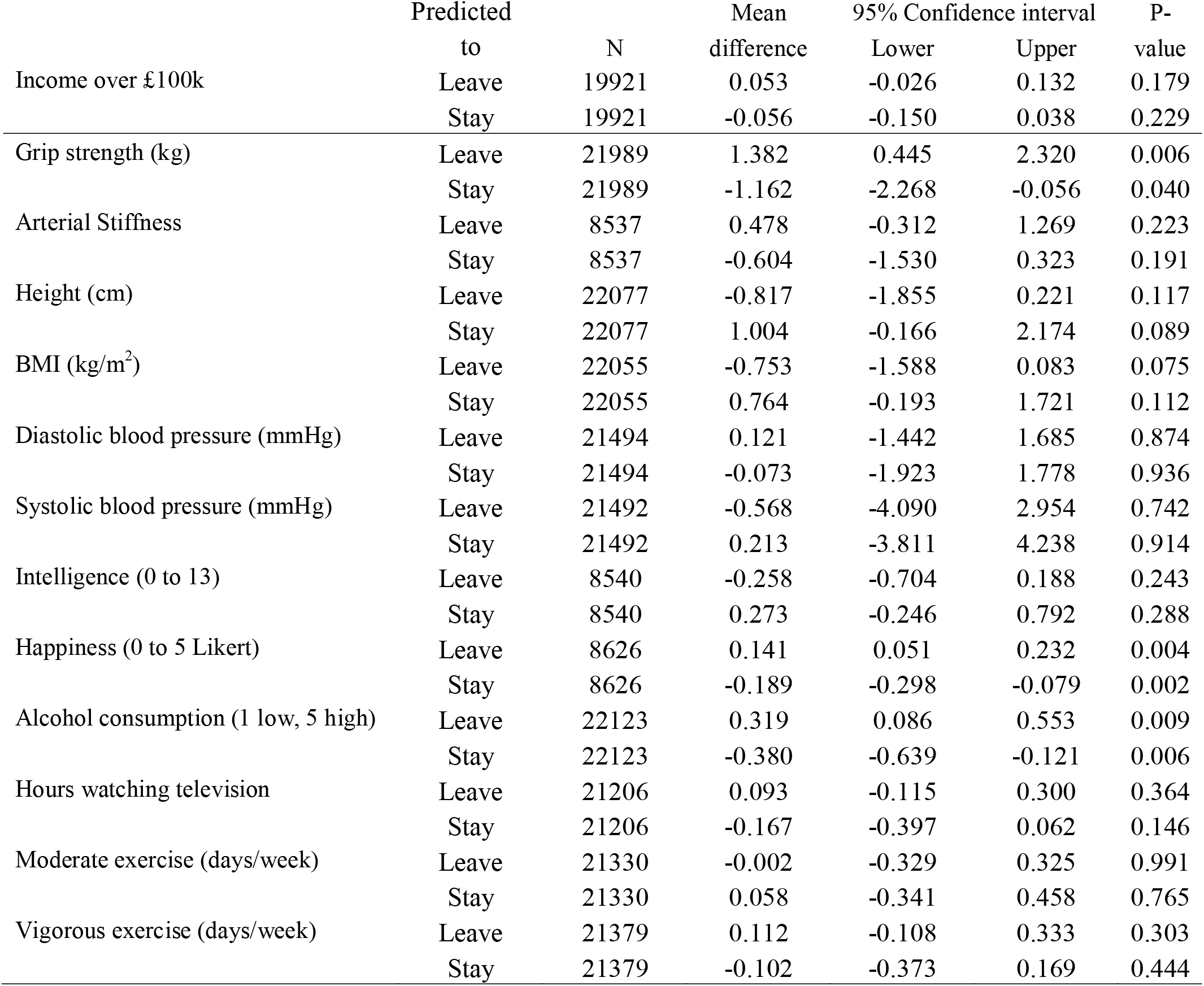
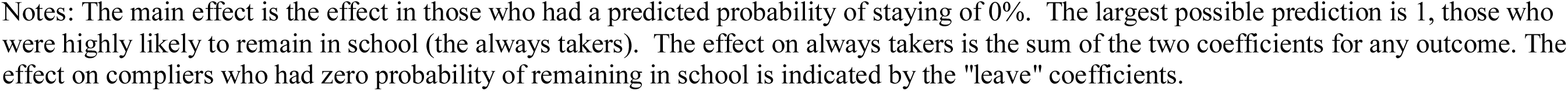
Heterogeneity in the effect of reform on outcomes by likelihood of staying in school. The effect of the reform on those who were predicted to leave at age 15 is indicated in the “leave” rows. The effect on those who were predicted to leave is the sum of the coefficients.

|  | Predicted<br>to | N | Mean<br>difference | 95% Confidence interval |  | P-value |
| --- | --- | --- | --- | --- | --- | --- |
|  |  |  |  | Lower | Upper |  |
| High blood pressure | Leave | 21768 | 0.049 | -0.014 | 0.113 | 0.123 |
|  | Stay | 21768 | -0.073 | -0.145 | -0.001 | 0.047 |
| Diabetes | Leave | 22049 | -0.034 | -0.060 | -0.009 | 0.011 |
|  | Stay | 22049 | 0.033 | 0.000 | 0.065 | 0.048 |
| Stroke | Leave | 22110 | 0.014 | 0.003 | 0.025 | 0.015 |
|  | Stay | 22110 | -0.020 | -0.033 | -0.007 | 0.004 |
| Heart attack | Leave | 22110 | -0.008 | -0.024 | 0.009 | 0.350 |
|  | Stay | 22110 | 0.007 | -0.013 | 0.028 | 0.473 |
| Depression | Leave | 21085 | -0.017 | -0.061 | 0.028 | 0.448 |
|  | Stay | 21085 | 0.013 | -0.040 | 0.066 | 0.612 |
| Cancer | Leave | 22011 | -0.010 | -0.037 | 0.017 | 0.439 |
|  | Stay | 22011 | 0.007 | -0.026 | 0.041 | 0.657 |
| Died | Leave | 22138 | 0.008 | -0.008 | 0.024 | 0.331 |
|  | Stay | 22138 | -0.014 | -0.032 | 0.004 | 0.124 |
| Ever smoked | Leave | 22086 | 0.058 | -0.009 | 0.126 | 0.087 |
|  | Stay | 22086 | -0.076 | -0.154 | 0.002 | 0.056 |
| Current smoker | Leave | 22086 | -0.032 | -0.095 | 0.031 | 0.308 |
|  | Stay | 22086 | 0.042 | -0.033 | 0.116 | 0.263 |
| Income over £18k | Leave | 19921 | -0.014 | -0.049 | 0.020 | 0.401 |
|  | Stay | 19921 | 0.013 | -0.034 | 0.060 | 0.560 |
| Income over £31k | Leave | 19921 | 0.081 | 0.003 | 0.159 | 0.043 |
|  | Stay | 19921 | -0.089 | -0.183 | 0.004 | 0.061 |
| Income over £52k | Leave | 19921 | 0.055 | -0.034 | 0.145 | 0.212 |
|  | Stay | 19921 | -0.039 | -0.143 | 0.065 | 0.446 |

|  | Predicted |  | Mean | 95% Confidence interval |  | P- |
| --- | --- | --- | --- | --- | --- | --- |
|  | to | N | difference | Lower | Upper | value |
| Income over £100k | Leave | 19921 | 0.053 | -0.026 | 0.132 | 0.179 |
|  | Stay | 19921 | -0.056 | -0.150 | 0.038 | 0.229 |
| Grip strength (kg) | Leave | 21989 | 1.382 | 0.445 | 2.320 | 0.006 |
|  | Stay | 21989 | -1.162 | -2.268 | -0.056 | 0.040 |
| Arterial Stiffness | Leave | 8537 | 0.478 | -0.312 | 1.269 | 0.223 |
|  | Stay | 8537 | -0.604 | -1.530 | 0.323 | 0.191 |
| Height (cm) | Leave | 22077 | -0.817 | -1.855 | 0.221 | 0.117 |
|  | Stay | 22077 | 1.004 | -0.166 | 2.174 | 0.089 |
| BMI (kg/m<sup>2</sup>) | Leave | 22055 | -0.753 | -1.588 | 0.083 | 0.075 |
|  | Stay | 22055 | 0.764 | -0.193 | 1.721 | 0.112 |
| Diastolic blood pressure (mmHg) | Leave | 21494 | 0.121 | -1.442 | 1.685 | 0.874 |
|  | Stay | 21494 | -0.073 | -1.923 | 1.778 | 0.936 |
| Systolic blood pressure (mmHg) | Leave | 21492 | -0.568 | -4.090 | 2.954 | 0.742 |
|  | Stay | 21492 | 0.213 | -3.811 | 4.238 | 0.914 |
| Intelligence (0 to 13) | Leave | 8540 | -0.258 | -0.704 | 0.188 | 0.243 |
|  | Stay | 8540 | 0.273 | -0.246 | 0.792 | 0.288 |
| Happiness (0 to 5 Likert) | Leave | 8626 | 0.141 | 0.051 | 0.232 | 0.004 |
|  | Stay | 8626 | -0.189 | -0.298 | -0.079 | 0.002 |
| Alcohol consumption (1 low, 5 high) | Leave | 22123 | 0.319 | 0.086 | 0.553 | 0.009 |
|  | Stay | 22123 | -0.380 | -0.639 | -0.121 | 0.006 |
| Hours watching television | Leave | 21206 | 0.093 | -0.115 | 0.300 | 0.364 |
|  | Stay | 21206 | -0.167 | -0.397 | 0.062 | 0.146 |
| Moderate exercise (days/week) | Leave | 21330 | -0.002 | -0.329 | 0.325 | 0.991 |
|  | Stay | 21330 | 0.058 | -0.341 | 0.458 | 0.765 |
| Vigorous exercise (days/week) | Leave | 21379 | 0.112 | -0.108 | 0.333 | 0.303 |
|  | Stay | 21379 | -0.102 | -0.373 | 0.169 | 0.444 |

Notes: The main effect is the effect in those who had a predicted probability of staying of 0%. The largest possible prediction is 1, those who were highly likely to remain in school (the always takers). The effect on always takers is the sum of the two coefficients for any outcome. The effect on compliers who had zero probability of remaining in school is indicated by the "leave" coefficients.

**Table S4:** Associations of outcome and the raising of the school leaving age in two negative control populations for MALES.

| Date of birth: | Negative control population 1 |  |  |  |  | Negative control population 2 |  |  |  |  |
| --- | --- | --- | --- | --- | --- | --- | --- | --- | --- | --- |
|  | Sept 1955-August 1956 to Sept 1956-August 1957 |  |  |  |  | Sept 1957- August 1958 to Sept 1958-August 1959 |  |  |  |  |
|  | N | Mean difference | 95% Confidence interval Lower | Upper | P-value | N | Mean difference | 95% Confidence interval Lower | Upper | P-value |
| High blood pressure* | 9,707 | -0.008 | -0.018 | 0.003 | 0.13 | 9,320 | -0.022 | -0.032 | -0.012 | <0.001 |
| Diabetes* | 9,790 | -0.002 | -0.008 | 0.003 | 0.40 | 9,388 | -0.001 | -0.006 | 0.004 | 0.61 |
| Stroke* | 9,834 | 0.000 | -0.002 | 0.002 | 0.90 | 9,430 | -0.003 | -0.006 | 0.000 | 0.08 |
| Heart attack* | 9,834 | -0.004 | -0.008 | -0.001 | 0.03 | 9,430 | 0.003 | -0.001 | 0.007 | 0.10 |
| Depression* | 9,526 | 0.007 | -0.005 | 0.019 | 0.22 | 9,072 | -0.003 | -0.015 | 0.009 | 0.60 |
| Cancer* | 9,834 | -0.004 | -0.012 | 0.003 | 0.25 | 9,406 | -0.005 | -0.013 | 0.002 | 0.17 |
| Died* | 9,852 | 0.002 | -0.002 | 0.005 | 0.35 | 9,439 | -0.002 | -0.005 | 0.001 | 0.23 |
| Ex-smoker* | 9,827 | -0.011 | -0.031 | 0.009 | 0.27 | 9,425 | 0.001 | -0.018 | 0.021 | 0.88 |
| Current smoker* | 9,827 | 0.000 | -0.012 | 0.012 | 0.99 | 9,425 | 0.004 | -0.010 | 0.018 | 0.57 |
| Income over £100k* | 9,047 | -0.004 | -0.013 | 0.005 | 0.35 | 8,628 | -0.001 | -0.009 | 0.008 | 0.88 |
| Income over £52k* | 9,047 | -0.003 | -0.019 | 0.013 | 0.70 | 8,628 | -0.007 | -0.023 | 0.008 | 0.35 |
| Income over £31k* | 9,047 | -0.007 | -0.018 | 0.005 | 0.23 | 8,628 | -0.014 | -0.023 | -0.004 | 0.006 |
| Income over £18k* | 9,047 | -0.003 | -0.014 | 0.007 | 0.52 | 8,628 | -0.008 | -0.017 | 0.001 | 0.07 |
| Grip strength (kg) | 9,773 | -0.158 | -0.446 | 0.129 | 0.27 | 9,368 | 0.058 | -0.323 | 0.440 | 0.75 |
| Arterial Stiffness | 3,746 | -0.229 | -0.407 | -0.051 | 0.01 | 3,564 | -0.168 | -0.318 | -0.017 | 0.03 |
| Height (cm) | 9,820 | 0.058 | -0.151 | 0.268 | 0.57 | 9,405 | 0.158 | 0.040 | 0.276 | 0.01 |
| BMI (kg/m<sup>2</sup>) | 9,804 | -0.139 | -0.288 | 0.011 | 0.07 | 9,399 | -0.012 | -0.157 | 0.133 | 0.86 |
| Diastolic blood pressure (mmHg) | 9,550 | -0.155 | -0.387 | 0.078 | 0.18 | 9,144 | 0.006 | -0.206 | 0.217 | 0.95 |
| Systolic blood pressure (mmHg) | 9,549 | -0.715 | -1.186 | -0.244 | 0.005 | 9,144 | -0.125 | -0.433 | 0.184 | 0.41 |
| Intelligence (0 to 13) | 3,710 | -0.063 | -0.148 | 0.021 | 0.14 | 3,542 | -0.009 | -0.150 | 0.132 | 0.90 |
| Happiness (0 to 5 Likert) | 3,757 | -0.026 | -0.055 | 0.003 | 0.08 | 3,574 | -0.012 | -0.043 | 0.019 | 0.43 |
| Alcohol consumption | 9,846 | -0.010 | -0.042 | 0.022 | 0.50 | 9,430 | -0.060 | -0.095 | -0.024 | 0.002 |
| Hours watching TV | 9,481 | 0.037 | -0.014 | 0.088 | 0.15 | 9,067 | -0.047 | -0.109 | 0.015 | 0.13 |
| Moderate exercise (days/wk) | 9,574 | 0.025 | -0.027 | 0.077 | 0.33 | 9,179 | -0.128 | -0.203 | -0.054 | 0.002 |
| Vigorous exercise (days/wk) | 9,554 | 0.037 | -0.023 | 0.097 | 0.22 | 9,172 | 0.038 | -0.010 | 0.087 | 0.12 |

Notes: \* denotes risk differences. Estimated using robust linear regression, with standard errors clustered by year and month of birth. All estimates adjust for month of birth and sex.

**Table S5:** Associations of outcomes and the raising of the school leaving age in two negative control populations <u>for FEMALES</u>.

| Date of birth: | Negative control population 1 |  |  |  |  | Negative control population 2 |  |  |  |  |
| --- | --- | --- | --- | --- | --- | --- | --- | --- | --- | --- |
|  | Sept 1955-August 1956 to Sept 1956-August 1957 |  |  |  |  | Sept 1957- August 1958 to Sept 1958-August 1959 |  |  |  |  |
|  | N | Mean difference | 95% Confidence interval |  | P-value | N | Mean difference | 95% Confidence interval |  | P-value |
| High blood pressure* | 12,697 | -0.012 | -0.022 | -0.002 | 0.02 | 11,833 | -0.009 | -0.019 | 0.001 | 0.07 |
| Diabetes* | 12,895 | 0.001 | -0.002 | 0.004 | 0.72 | 11,988 | -0.002 | -0.005 | 0.001 | 0.14 |
| Stroke* | 12,922 | 0.001 | -0.001 | 0.002 | 0.33 | 12,022 | 0.001 | -0.002 | 0.003 | 0.63 |
| Heart attack* | 12,922 | 0.000 | -0.002 | 0.001 | 0.44 | 12,022 | 0.001 | -0.001 | 0.002 | 0.45 |
| Depression* | 12,132 | 0.015 | 0.005 | 0.025 | 0.007 | 11,356 | -0.004 | -0.014 | 0.005 | 0.37 |
| Cancer* | 12,836 | -0.006 | -0.013 | 0.001 | 0.09 | 11,928 | -0.008 | -0.015 | 0.000 | 0.05 |
| Died* | 12,937 | 0.002 | -0.001 | 0.004 | 0.14 | 12,037 | 0.003 | 0.001 | 0.004 | <0.001 |
| Ex-smoker* | 12,901 | -0.011 | -0.019 | -0.003 | 0.008 | 12,011 | -0.003 | -0.016 | 0.010 | 0.68 |
| Current smoker* | 12,901 | 0.000 | -0.009 | 0.009 | 0.99 | 12,011 | 0.004 | -0.002 | 0.010 | 0.22 |
| Income over £100k* | 11,384 | 0.010 | 0.002 | 0.019 | 0.02 | 10,738 | 0.006 | 0.000 | 0.013 | 0.06 |
| Income over £52k* | 11,384 | 0.010 | -0.004 | 0.024 | 0.15 | 10,738 | 0.014 | 0.001 | 0.027 | 0.03 |
| Income over £31k* | 11,384 | 0.007 | -0.007 | 0.020 | 0.30 | 10,738 | 0.017 | 0.007 | 0.027 | 0.002 |
| Income over £18k* | 11,384 | 0.014 | 0.009 | 0.020 | <0.001 | 10,738 | 0.009 | -0.001 | 0.019 | 0.09 |
| Grip strength (kg) | 12,862 | 0.355 | 0.188 | 0.523 | <0.001 | 11,968 | 0.187 | 0.072 | 0.301 | 0.003 |
| Arterial Stiffness | 5,018 | -0.248 | -0.356 | -0.140 | <0.001 | 4,589 | -0.204 | -0.313 | -0.095 | <0.001 |
| Height (cm) | 12,907 | 0.101 | -0.054 | 0.257 | 0.19 | 12,017 | 0.159 | 0.008 | 0.310 | 0.04 |
| BMI (kg/m<sup>2</sup>) | 12,889 | 0.049 | -0.071 | 0.168 | 0.41 | 12,012 | -0.069 | -0.196 | 0.058 | 0.27 |
| Diastolic blood pressure (mmHg) | 12,551 | -0.390 | -0.756 | -0.023 | 0.04 | 11,685 | -0.565 | -0.887 | -0.243 | 0.001 |
| Systolic blood pressure (mmHg) | 12,550 | -1.047 | -1.586 | -0.508 | <0.001 | 11,685 | -1.044 | -1.448 | -0.640 | <0.001 |
| Intelligence (0 to 13) | 5,045 | -0.086 | -0.167 | -0.004 | 0.04 | 4,621 | -0.102 | -0.159 | -0.045 | 0.001 |
| Happiness (0 to 5 Likert) | 5,090 | 0.033 | 0.000 | 0.066 | 0.05 | 4,653 | -0.004 | -0.024 | 0.015 | 0.66 |
| Alcohol consumption | 12,932 | -0.013 | -0.054 | 0.028 | 0.53 | 12,031 | -0.006 | -0.043 | 0.031 | 0.74 |
| Hours watching TV | 12,387 | -0.028 | -0.075 | 0.019 | 0.23 | 11,492 | -0.060 | -0.100 | -0.021 | 0.004 |
| Moderate exercise (days/wk) | 12,384 | 0.014 | -0.059 | 0.086 | 0.70 | 11,550 | 0.028 | -0.053 | 0.108 | 0.48 |
| Vigorous exercise (days/wk) | 12,440 | 0.082 | 0.033 | 0.132 | 0.002 | 11,658 | 0.089 | 0.036 | 0.141 | 0.002 |

Notes: \* denotes risk differences. Estimated using robust linear regression, with standard errors clustered by year and month of birth.

**Table S6:**
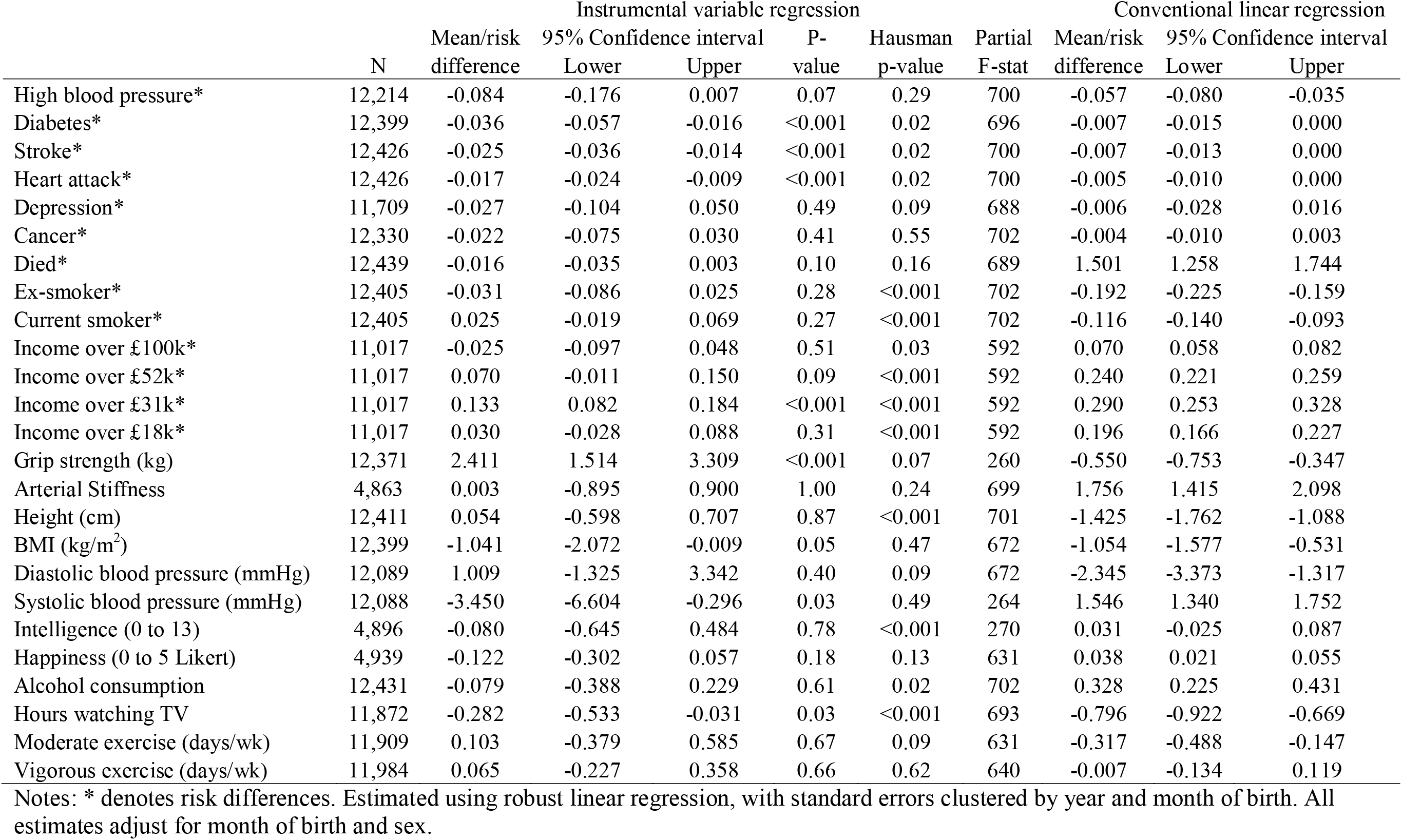
The effects of leaving school after age 15, instrumental variable regression (left) and conventional regression (right), <u>MALES</u>.

**Table S7:**
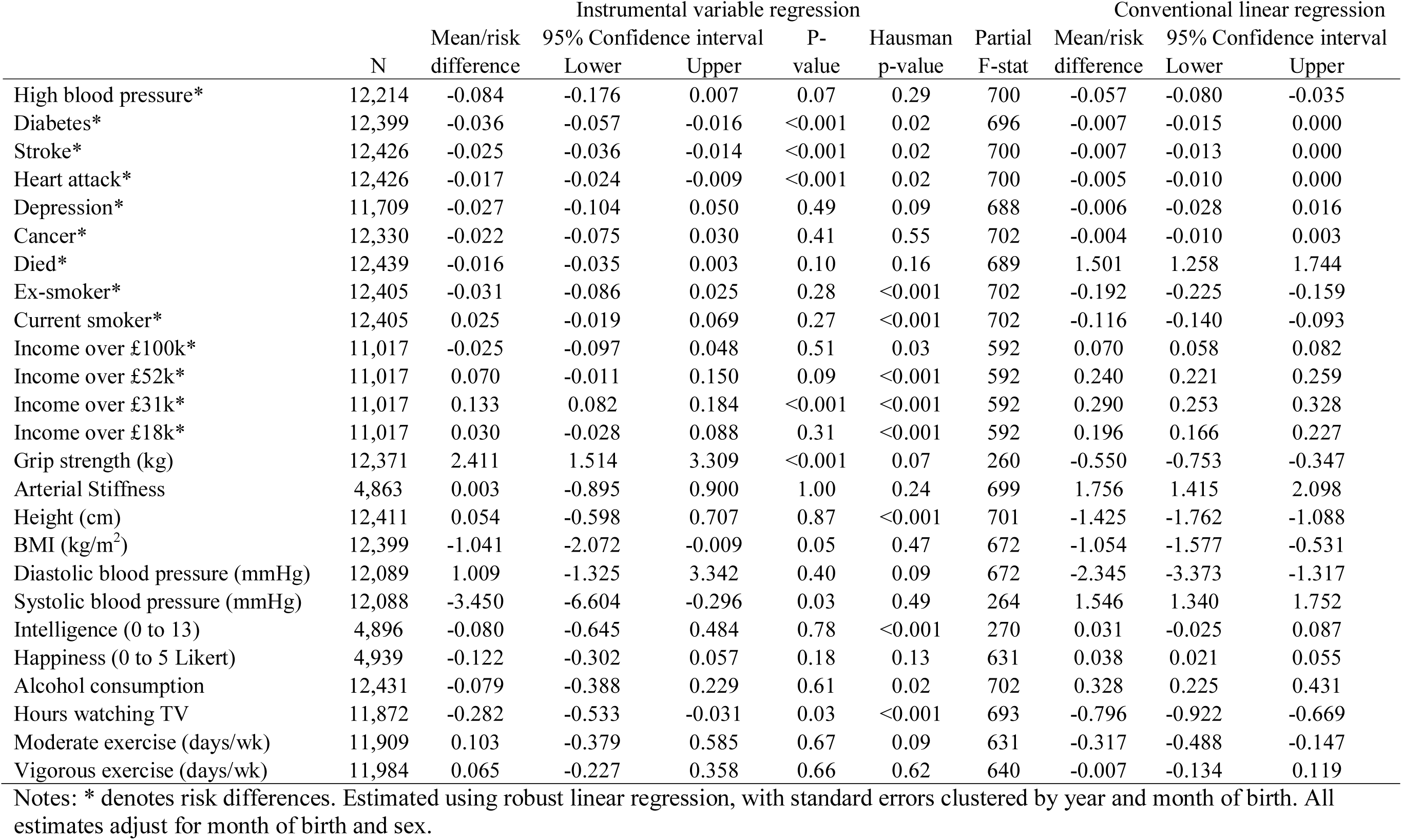
The effects of leaving school after age 15, instrumental variable regression (left) and conventional regression (right), <u>FEMALES</u>.

**Table S8:** The effects of leaving school after age 15, instrumental variable regression (left) and conventional regression (right), Clark and Royer (2013) specification 47 month bandwidth, <u>MALES and FEMALES</u>.

|  | N | Instrumental variable regression |  |  |  | Conventional linear regression |  |  |  |  |
| --- | --- | --- | --- | --- | --- | --- | --- | --- | --- | --- |
|  |  | Mean/risk difference | 95% Confidence interval Lower | 95% Confidence interval Upper | P-value | Hausman p-value | Partial F-stat | Mean/risk difference | 95% Confidence interval Lower | 95% Confidence interval Upper |
| High blood pressure* | 288,579 | -0.069 | -0.235 | 0.096 | 0.41 | 0.75 | 75 | -0.043 | -0.047 | -0.039 |
| Diabetes* | 295,606 | 0.073 | 0.001 | 0.146 | 0.05 | 0.04 | 81 | -0.018 | -0.020 | -0.016 |
| Stroke* | 296,729 | 0.062 | 0.015 | 0.109 | 0.009 | 0.02 | 80 | -0.009 | -0.011 | -0.008 |
| Heart attack* | 296,729 | 0.172 | 0.117 | 0.226 | <0.001 | 1.2E-4 | 80 | -0.018 | -0.020 | -0.016 |
| Depression* | 281,588 | -2.385 | -3.252 | -1.519 | <0.001 | 1.6E-5 | 66 | 0.032 | 0.029 | 0.036 |
| Cancer* | 296,144 | 0.237 | 0.124 | 0.350 | <0.001 | 0.001 | 79 | 0.002 | -0.001 | 0.006 |
| Died* | 297,226 | 0.147 | 0.090 | 0.205 | <0.001 | 0.001 | 80 | -0.007 | -0.008 | -0.006 |
| Ex-smoker* | 296,106 | -1.148 | -1.489 | -0.808 | <0.001 | 2.4E-5 | 78 | -0.102 | -0.107 | -0.097 |
| Current smoker* | 296,106 | 0.120 | -0.029 | 0.268 | 0.11 | 0.04 | 78 | -0.054 | -0.057 | -0.050 |
| Income over £100k* | 251,968 | -1.529 | -1.889 | -1.170 | <0.001 | 3.3E-6 | 64 | 0.285 | 0.275 | 0.294 |
| Income over £52k* | 251,968 | -1.432 | -1.837 | -1.027 | <0.001 | 2.0E-6 | 64 | 0.270 | 0.265 | 0.275 |
| Income over £31k* | 251,968 | -0.675 | -1.017 | -0.333 | <0.001 | 5.5E-5 | 64 | 0.151 | 0.144 | 0.158 |
| Income over £18k* | 251,968 | -0.023 | -0.159 | 0.113 | 0.74 | 0.42 | 64 | 0.033 | 0.031 | 0.035 |
| Grip strength (kg) | 294,729 | 3.140 | -1.928 | 8.208 | 0.22 | 0.44 | 80 | 1.214 | 1.150 | 1.278 |
| Arterial Stiffness | 116,704 | -7.251 | -10.474 | -4.028 | <0.001 | 4.2E-4 | 23 | -0.211 | -0.266 | -0.156 |
| Height (cm) | 296,227 | -5.085 | -8.861 | -1.309 | 0.008 | 0.007 | 80 | 2.100 | 2.041 | 2.159 |
| BMI (kg/m<sup>2</sup>) | 295,900 | -1.702 | -3.465 | 0.060 | 0.06 | 0.56 | 80 | -1.174 | -1.219 | -1.129 |
| Diastolic blood pressure (mmHg) | 288,385 | -10.775 | -16.293 | -5.256 | <0.001 | 0.003 | 75 | -0.322 | -0.414 | -0.229 |
| Systolic blood pressure (mmHg) | 288,383 | -7.456 | -14.956 | 0.043 | 0.05 | 0.14 | 75 | -1.432 | -1.605 | -1.259 |
| Intelligence (0 to 13) | 115,497 | -13.829 | -18.127 | -9.531 | <0.001 | 4.6E-7 | 27 | 1.761 | 1.731 | 1.790 |
| Happiness (0 to 5 Likert) | 117,859 | -0.562 | -1.249 | 0.124 | 0.11 | 0.11 | 25 | -0.011 | -0.022 | -0.001 |
| Alcohol consumption | 297,051 | -1.473 | -2.357 | -0.590 | 0.001 | 0.001 | 80 | 0.476 | 0.463 | 0.489 |
| Hours watching TV | 286,794 | 1.299 | 0.580 | 2.018 | <0.001 | 1.4E-4 | 85 | -0.833 | -0.849 | -0.816 |
| Moderate exercise (days/wk) | 282,844 | 1.823 | 0.531 | 3.115 | 0.006 | 0.002 | 68 | -0.270 | -0.295 | -0.246 |
| Vigorous exercise (days/wk) | 282,863 | 2.413 | 1.097 | 3.729 | <0.001 | 1.6E-4 | 69 | -0.123 | -0.142 | -0.103 |

Notes: \* denotes risk differences. Estimated using robust linear regression, with standard errors clustered by year and month of birth. All estimates adjust for month of birth, sex, and linear time-trend.

**Table S9:** The effects of leaving school after age 15, instrumental variable regression (left) and conventional regression (right), Clark and Royer (2013) specification 47 month bandwidth, <u>MALES</u>.

|  | N | Instrumental variable regression |  |  |  |  | Partial F-stat | Conventional linear regression |  |  |
| --- | --- | --- | --- | --- | --- | --- | --- | --- | --- | --- |
|  |  | Mean/risk difference | 95% Confidence interval Lower | Upper | P-value | Hausman p-value |  | Mean/risk difference | 95% Confidence interval Lower | Upper |
| High blood pressure* | 135,775 | -0.376 | -0.710 | -0.043 | 0.03 | 0.04 | 23 | -0.045 | -0.051 | -0.038 |
| Diabetes* | 137,554 | 0.028 | -0.113 | 0.170 | 0.69 | 0.51 | 24 | -0.020 | -0.024 | -0.017 |
| Stroke* | 138,282 | 0.065 | -0.019 | 0.149 | 0.13 | 0.10 | 24 | -0.011 | -0.013 | -0.008 |
| Heart attack* | 138,282 | 0.231 | 0.097 | 0.364 | <0.001 | 0.003 | 24 | -0.030 | -0.033 | -0.027 |
| Depression* | 133,061 | -2.570 | -3.596 | -1.545 | <0.001 | 1.0E-4 | 20 | 0.028 | 0.024 | 0.033 |
| Cancer* | 138,320 | 0.542 | 0.328 | 0.756 | <0.001 | 1.4E-4 | 24 | 0.003 | -0.002 | 0.007 |
| Died* | 138,507 | 0.318 | 0.182 | 0.454 | <0.001 | 5.5E-5 | 24 | -0.012 | -0.014 | -0.010 |
| Ex-smoker* | 137,960 | -1.191 | -1.797 | -0.586 | <0.001 | 0.004 | 23 | -0.107 | -0.114 | -0.101 |
| Current smoker* | 137,960 | 0.080 | -0.237 | 0.397 | 0.62 | 0.43 | 23 | -0.053 | -0.058 | -0.048 |
| Income over £100k* | 123,137 | -1.832 | -2.491 | -1.173 | <0.001 | 5.7E-7 | 22 | 0.291 | 0.278 | 0.304 |
| Income over £52k* | 123,137 | -2.545 | -3.449 | -1.640 | <0.001 | 6.2E-7 | 22 | 0.303 | 0.296 | 0.310 |
| Income over £31k* | 123,137 | -1.577 | -2.293 | -0.862 | <0.001 | 3.8E-5 | 22 | 0.181 | 0.173 | 0.190 |
| Income over £18k* | 123,137 | -0.277 | -0.533 | -0.021 | 0.03 | 0.01 | 22 | 0.042 | 0.039 | 0.045 |
| Grip strength (kg) | 137,365 | -8.106 | -17.672 | 1.460 | 0.10 | 0.09 | 25 | 0.973 | 0.862 | 1.084 |
| Arterial Stiffness | 55,196 | -13.426 | -23.482 | -3.369 | 0.009 | 0.001 | 6 | -0.218 | -0.317 | -0.118 |
| Height (cm) | 137,989 | -10.128 | -15.884 | -4.372 | <0.001 | 6.3E-4 | 25 | 2.279 | 2.194 | 2.365 |
| BMI (kg/m<sup>2</sup>) | 137,835 | -2.824 | -6.485 | 0.838 | 0.13 | 0.36 | 25 | -1.056 | -1.114 | -0.998 |
| Diastolic blood pressure (mmHg) | 134,450 | -12.780 | -24.648 | -0.913 | 0.03 | 0.05 | 24 | -0.315 | -0.443 | -0.187 |
| Systolic blood pressure (mmHg) | 134,449 | -11.376 | -26.293 | 3.542 | 0.14 | 0.18 | 24 | -1.329 | -1.566 | -1.092 |
| Intelligence (0 to 13) | 54,397 | -21.791 | -34.286 | -9.297 | <0.001 | 1.2E-7 | 7 | 1.858 | 1.818 | 1.899 |
| Happiness (0 to 5 Likert) | 55,574 | -0.954 | -2.371 | 0.464 | 0.19 | 0.19 | 6 | -0.025 | -0.039 | -0.010 |
| Alcohol consumption | 138,428 | -3.425 | -4.992 | -1.857 | <0.001 | 1.9E-4 | 24 | 0.407 | 0.388 | 0.425 |
| Hours watching TV | 133,969 | 3.613 | 1.935 | 5.292 | <0.001 | 1.8E-5 | 25 | -0.833 | -0.857 | -0.808 |
| Moderate exercise (days/wk) | 133,519 | 4.371 | 1.663 | 7.078 | 0.002 | 3.3E-4 | 23 | -0.476 | -0.511 | -0.441 |
| Vigorous exercise (days/wk) | 133,008 | 3.844 | 1.178 | 6.510 | 0.005 | 0.008 | 22 | -0.243 | -0.271 | -0.215 |

Notes: \* denotes risk differences. Estimated using robust linear regression, with standard errors clustered by year and month of birth. All estimates adjust for month of birth, sex, and linear time-trend.

**Table S10:** The effects of leaving school after age 15, instrumental variable regression (left) and conventional regression (right), Clark and Royer (2013) specification 47 month bandwidth, <u>FEMALES</u>.

|  | N | Instrumental variable regression |  |  |  |  | Conventional linear regression |  |  |  |
| --- | --- | --- | --- | --- | --- | --- | --- | --- | --- | --- |
|  |  | Mean/risk difference | 95% Confidence interval |  | P-value | Hausman p-value | Partial F-stat | Mean/risk difference | 95% Confidence interval |  |
|  |  |  | Lower | Upper |  |  |  |  | Lower | Upper |
| High blood pressure* | 152,804 | 0.088 | -0.065 | 0.242 | 0.26 | 0.12 | 54 | -0.041 | -0.046 | -0.036 |
| Diabetes* | 158,052 | 0.074 | -0.002 | 0.149 | 0.06 | 0.06 | 58 | -0.016 | -0.018 | -0.013 |
| Stroke* | 158,447 | 0.051 | 0.000 | 0.101 | 0.05 | 0.06 | 58 | -0.008 | -0.010 | -0.007 |
| Heart attack* | 158,447 | 0.104 | 0.066 | 0.142 | <0.001 | 4.6E-5 | 58 | -0.008 | -0.009 | -0.006 |
| Depression* | 148,527 | -2.326 | -3.283 | -1.368 | <0.001 | 1.1E-5 | 47 | 0.036 | 0.032 | 0.040 |
| Cancer* | 157,824 | 0.025 | -0.105 | 0.156 | 0.70 | 0.73 | 56 | 0.002 | -0.002 | 0.007 |
| Died* | 158,719 | 0.031 | -0.029 | 0.090 | 0.31 | 0.32 | 57 | -0.003 | -0.004 | -0.001 |
| Ex-smoker* | 158,146 | -1.232 | -1.602 | -0.862 | <0.001 | 1.6E-6 | 57 | -0.097 | -0.104 | -0.091 |
| Current smoker* | 158,146 | 0.150 | 0.004 | 0.297 | 0.04 | 0.01 | 57 | -0.054 | -0.058 | -0.050 |
| Income over £100k* | 128,831 | -1.410 | -1.818 | -1.002 | <0.001 | 3.9E-5 | 45 | 0.279 | 0.271 | 0.287 |
| Income over £52k* | 128,831 | -0.757 | -1.131 | -0.383 | <0.001 | 1.6E-4 | 45 | 0.236 | 0.229 | 0.244 |
| Income over £31k* | 128,831 | -0.069 | -0.402 | 0.265 | 0.69 | 0.25 | 45 | 0.120 | 0.112 | 0.128 |
| Income over £18k* | 128,831 | 0.150 | 0.015 | 0.285 | 0.03 | 0.07 | 45 | 0.024 | 0.022 | 0.027 |
| Grip strength (kg) | 157,364 | 10.449 | 6.021 | 14.877 | <0.001 | 4.0E-5 | 57 | 1.423 | 1.354 | 1.492 |
| Arterial Stiffness | 61,508 | -3.879 | -6.866 | -0.892 | 0.01 | 0.03 | 18 | -0.205 | -0.272 | -0.138 |
| Height (cm) | 158,238 | -2.218 | -6.422 | 1.987 | 0.30 | 0.10 | 57 | 1.940 | 1.863 | 2.017 |
| BMI (kg/m<sup>2</sup>) | 158,065 | -0.590 | -3.229 | 2.049 | 0.66 | 0.61 | 56 | -1.277 | -1.339 | -1.215 |
| Diastolic blood pressure (mmHg) | 153,935 | -8.062 | -12.685 | -3.438 | <0.001 | 0.01 | 53 | -0.327 | -0.448 | -0.205 |
| Systolic blood pressure (mmHg) | 153,934 | 0.539 | -8.079 | 9.158 | 0.90 | 0.64 | 53 | -1.500 | -1.735 | -1.264 |
| Intelligence (0 to 13) | 61,100 | -9.694 | -13.101 | -6.287 | <0.001 | 1.4E-5 | 21 | 1.671 | 1.634 | 1.708 |
| Happiness (0 to 5 Likert) | 62,285 | -0.434 | -1.203 | 0.334 | 0.27 | 0.24 | 20 | 0.000 | -0.013 | 0.014 |
| Alcohol consumption | 158,623 | -0.585 | -1.527 | 0.357 | 0.22 | 0.03 | 58 | 0.536 | 0.518 | 0.554 |
| Hours watching TV | 152,825 | 0.121 | -0.995 | 1.237 | 0.83 | 0.13 | 61 | -0.832 | -0.852 | -0.811 |
| Moderate exercise (days/wk) | 149,325 | 0.376 | -0.998 | 1.751 | 0.59 | 0.51 | 47 | -0.082 | -0.111 | -0.053 |
| Vigorous exercise (days/wk) | 149,855 | 1.753 | 0.323 | 3.183 | 0.02 | 0.004 | 49 | -0.013 | -0.038 | 0.012 |

Notes: \* denotes risk differences. Estimated using robust linear regression, with standard errors clustered by year and month of birth. All estimates adjust for month of birth, sex, and linear time-trend.

